# Replicative selfish genetic elements are driving rapid pathogenic adaptation of *Enterococcus faecium*

**DOI:** 10.1101/2025.03.16.643550

**Authors:** Matthew P Grieshop, Aaron A Behr, Sierra Bowden, Jordan D Lin, Marco Molari, Gabriella ZM Reynolds, Erin F Brooks, Boryana Doyle, Guillermo Rodriguez-Nava, Jorge L Salinas, Niaz Banaei, Ami S Bhatt

## Abstract

Understanding how healthcare-associated pathogens adapt in clinical environments can inform strategies to reduce their burden. Here, we investigate the hypothesis that insertion sequences (IS), prokaryotic transposable elements, are a dominant mediator of rapid genomic evolution in healthcare-associated pathogens. Among 28,207 publicly available pathogen genomes, we find high copy numbers of the replicative ISL3 family in healthcare-associated *Enterococcus faecium, Streptococcus pneumoniae and Staphylococcus aureus*. In *E. faecium,* the ESKAPE pathogen with the highest IS density, we find that ISL3 proliferation has increased in the last 30 years. To enable better identification of structural variants, we long read-sequenced a new, single hospital collection of 282 *Enterococcal* infection isolates collected over three years. In these samples, we observed extensive, ongoing structural variation of the *E. faecium* genome, largely mediated by active replicative ISL3 elements. To determine if ISL3 is actively replicating in clinical timescales in its natural, gut microbiome reservoir, we long read-sequenced a collection of 28 longitudinal stool samples from patients undergoing hematopoietic cell transplantation, whose gut microbiomes were dominated by *E. faecium*. We found up to six structural variants of a given *E. faecium* strain within a single stool sample. Examining longitudinal samples from one individual in further detail, we find ISL3 elements can replicate and move to specific positions with profound regulatory effects on neighboring gene expression. In particular, we identify an ISL3 element that upon insertion replaces an imperfect -35 promoter sequence at a *folT* gene locus with a perfect -35 sequence, which leads to substantial upregulation of expression of *folT*, driving highly effective folate scavenging. As a known folate auxotroph, *E. faecium* depends on other members of the microbiota or diet to supply folate. Enhanced folate scavenging may enable *E. faecium* to thrive in the setting of microbiome collapse that is common in HCT and other critically ill patients. Together, ISL3 expansion has enabled *E. faecium* to rapidly evolve in healthcare settings, and this likely contributes to its metabolic fitness and may strongly influence its ongoing trajectory of genomic evolution.

## Introduction

Healthcare-associated bacterial pathogens persist in the face of diverse and extreme selective pressures. Amidst repeated population bottlenecks, they navigate antibiotic treatments, antiseptics, off-target effects of human drugs, host immune defenses and fluctuating nutrient availability to colonize surfaces and patients within the hospital environment. The ability to rapidly adapt under these conditions is a defining feature of successful healthcare-associated pathogens. While numerous studies of pathogen evolution have revealed single-nucleotide polymorphisms (SNPs) and small insertions and deletions (indels) contributing to clinical adaptation^1–4^, the role of larger genomic structural variation mediated by selfish genetic elements – those promoting their own propagation within a genome without a direct fitness benefit – is understudied. Such elements can drive gain and loss of functional processes that enable bacteria to escape phage predation^5^, resist antibiotics^6,7^, and improve metabolic efficiency^8^ in laboratory conditions. However, these controlled environments may not fully capture the complexity of adaptation in clinical settings. Early comparative genomics analyses of bacterial pathogen genomes, carried out during the era of ‘complete, closed genome’ generation through the use of Sanger sequencing, elucidated examples of structural variation over long evolutionary timescales^9,10^. Yet, because recent large-scale genomic studies of clinical isolates primarily use short-read sequencing technologies, which often fail to resolve repetitive transposable elements associated with structural variations, our understanding of these changes during short-term clinical adaptation remains limited^1,4,11^. This narrow perspective highlights a significant gap in our knowledge of how pathogens continue to evolve in clinical environments.

Selfish transposable elements, particularly insertion sequences (IS), are well positioned to drive rapid evolutionary change in bacterial pathogens. These genetic elements are ubiquitous across the tree of life, with transposases being the most abundant genes in nature^12,13^. IS elements, the simplest autonomous transposable elements, typically carry only a transposase gene and employ diverse chemistries for either conservative ‘cut-and-paste’ or replicative ‘copy-and-paste’ mechanisms^13^. On long timescales (hundreds to millions of years), IS elements can drive speciation processes^9,10^. Briefly, IS elements expand within a genome, then recombination between copies and loss of the intervening sequence leads to genome reduction, as seen in the evolution of host-restricted pathogens from avirulent ancestors, such as in the case of *Bordetella pertussis*, *Yersinia pestis* and Shigella species^9,10,14^. Organisms that lack CRISPR systems or other defense systems against transposable element activity, such as *Enterococcus faecium*^15^, have been described to accumulate various mobile genetic elements^16,17^. To date, subsequent genome reduction in *E. faecium* has not been reported.

Transposable elements are also known for their ability to induce structural variation on short timescales and at relatively high rates^5,18^. Experimental evolution studies in bacteria have shown that IS-mediated mutations arise at rates many orders of magnitude greater than SNPs and indels^5,19^ and are responsible for the majority of the mutation burden when population size is small^19^. These variants can have profound effects on bacterial genomes. IS can insert into and disrupt genes^13^, while intergenic insertions can generate hybrid promoters^20^, leading to altered gene expression. Additionally, repetitive IS elements can serve as substrates for recombination leading to larger scale deletions and inversions^13,21^. This has been demonstrated by early, parallel mutations in the *Escherichia coli* long-term evolution experiment (LTEE) revealing IS-mediated deletion of the ribose operon sweeping through all populations within weeks^8,22^.

The consequences of IS-mediated genomic plasticity are manyfold. In clinical settings, such flexibility could potentially facilitate rapid adaptation to antibiotic pressures, immune responses, or new ecological niches within the host. Yet, the extent and ways in which IS elements actively shape bacterial genomes within natural populations on clinical (days to weeks) timescales remains an open question. Given frequent population bottlenecks, small population sizes in hosts or on hospital surfaces, and various strong selective pressures, we hypothesize that IS elements, especially those that are replicative and thus can increase in copy number within a genome, also mediate structural variants contributing to the evolution of bacterial pathogens on clinical timescales. Understanding how these mechanisms operate on short timescales in clinical isolates could provide crucial insights into the ongoing evolution of healthcare-associated pathogens and inform strategies to combat their persistence and spread.

The ESKAPE pathogens (*Enterobacter spp.*, *Staphylococcus aureus*, *Klebsiella pneumoniae, Acinetobacter baumannii, Pseudomonas aeruginosa, Enterococcus faecium/faecalis* and *Escherichia coli*) are disproportionately responsible for healthcare-associated infections^23^. These pathogens are ideal systems for studying the role of IS elements in clinical adaptation due to their success in the hospital environment and the wealth of available recent genomic data. Although no comprehensive study has contextualized the extent or role of transposable elements across these taxa, some reports have associated their pathogenicity with an enrichment of IS elements^24–28^. For example, prior studies of *E. faecium* and *E. faecalis*^28,29^, two gut-dwelling bacteria that can be pathogenic when found in the bloodstream, wounds, or abscesses, identified the signatures of IS expansion in the emergence of healthcare-associated lineages. Specifically, the presence of the replicative IS16 element of the IS256 family distinguishes healthcare-associated lineages of *E. faecium* from commensal ones, implicating IS16 elements in *E. faecium*’s pathogenicity^29^. However, a comprehensive understanding of how replicative IS elements contribute to the active evolution of pathogens, at either the population level or within individual patients, is lacking.

In this study, we investigate the role of IS elements in the rapid evolution of healthcare-associated pathogens. We report a large-scale survey of IS elements in complete pathogen genomes which reveals that while IS elements (both non-replicative and replicative) are abundant across healthcare-associated pathogens, *E. faecium* is distinguished by the highest IS density. In contrast to previous reports^17,29^, we identify ISL3 family elements as the predominant replicative IS in *E. faecium* genomes. Retrospective analysis of hospital-associated *E. faecium* populations since 1990 demonstrates that ISL3 elements have undergone marked expansion, coinciding with *E. faecium*’s increasing clinical importance^30,31^. To characterize the extent and impact of IS-mediated structural variation, we employ long-read DNA sequencing of clinical *E. faecium* isolates from a modern hospital-endemic lineage. This approach reveals that ISL3 elements contribute disproportionately to extensive recent genome rearrangements. Because *E. faecium* colonizes and often dominates the gastrointestinal tract of critically ill patients, examining longitudinal stool samples from these individuals presents a unique opportunity to study bacterial adaptation in real-time. To this end, we conducted longitudinal metagenomic analyses of gut microbiome samples from individuals undergoing allogeneic hematopoietic cell transplant (HCT) with high relative abundance of *E. faecium*. This approach revealed dynamic population heterogeneity and IS-mediated structural variations within individual patients over the course of their treatment. Subsequent isolation of these variants and RNA-seq analysis demonstrated the evolution of variants with different metabolic behavior, driven by an ISL3 insertion. Notably, an ISL3 insertion upstream of *folT* generated a strong hybrid promoter substantially enhancing folate scavenging, a trait that appears to confer a significant advantage in the gut environment of critically ill patients. Taken together, careful quantification and placement of selfish genetic elements in bacterial pathogen genomes reveals an active, ISL3-dependent mechanism for *E. faecium* evolution and proposes a path for its gut domination, eventual genome reduction and hospital-restriction.

## Results

### Insertion Sequences across the ESKAPE pathogens

Early surveys in prokaryotes found high IS content heterogeneity, even within closely related species^25,26^. However, this analysis was limited by the number of available genomes. The increasing availability of complete genomes, enabled by the transition toward long-read sequencing from short-read sequencing, now allows the study of thousands of high-contiguity pathogen genomes, presenting an opportunity to systematically map and directly compare IS distributions. Such comparisons are crucial for understanding the relative significance of IS elements in different pathogens and their relevance to pathogens in clinical environments.

To systematically characterize the distribution and diversity of IS elements across clinically important bacteria, we quantified IS content in 20,377 highly contiguous genomes from the ESKAPE pathogens: ***E****nterococcus faecium* (639) and *faecalis* (913), ***S****taphylococcus aureus* (2672)*, **K**lebsiella pneumoniae* (4627)*, **A**cinetobacter baumannii* (1082)*, **P**seudomonas aeruginosa* (1399)*, **E**nterobacter spp.* (526), and ***E****scherichia coli* (8519) (Fig. 1A). For comparison, we also analyzed 7,830 genomes from other species in the ESKAPE genera as well as several other taxa of clinical significance, and/or with notable IS expansions (*Shigella*, *Bordetella*, *Streptococcus*, *Salmonella,* Extended Data Fig. 1). We used both publicly available Oxford Nanopore Technologies (ONT) isolate genomic data and ‘complete’ assemblies from these taxa. To standardize processing of long-read sequencing data, we developed a workflow consisting of *de novo* assembly, genome quality control, and taxonomic verification, and then identified IS elements in each assembly using ISEScan^32^ (Supplementary Data 1). As IS elements are often carried on plasmids, we distinguished chromosomal and plasmid contigs with geNomad^33^.

**Figure 1.**
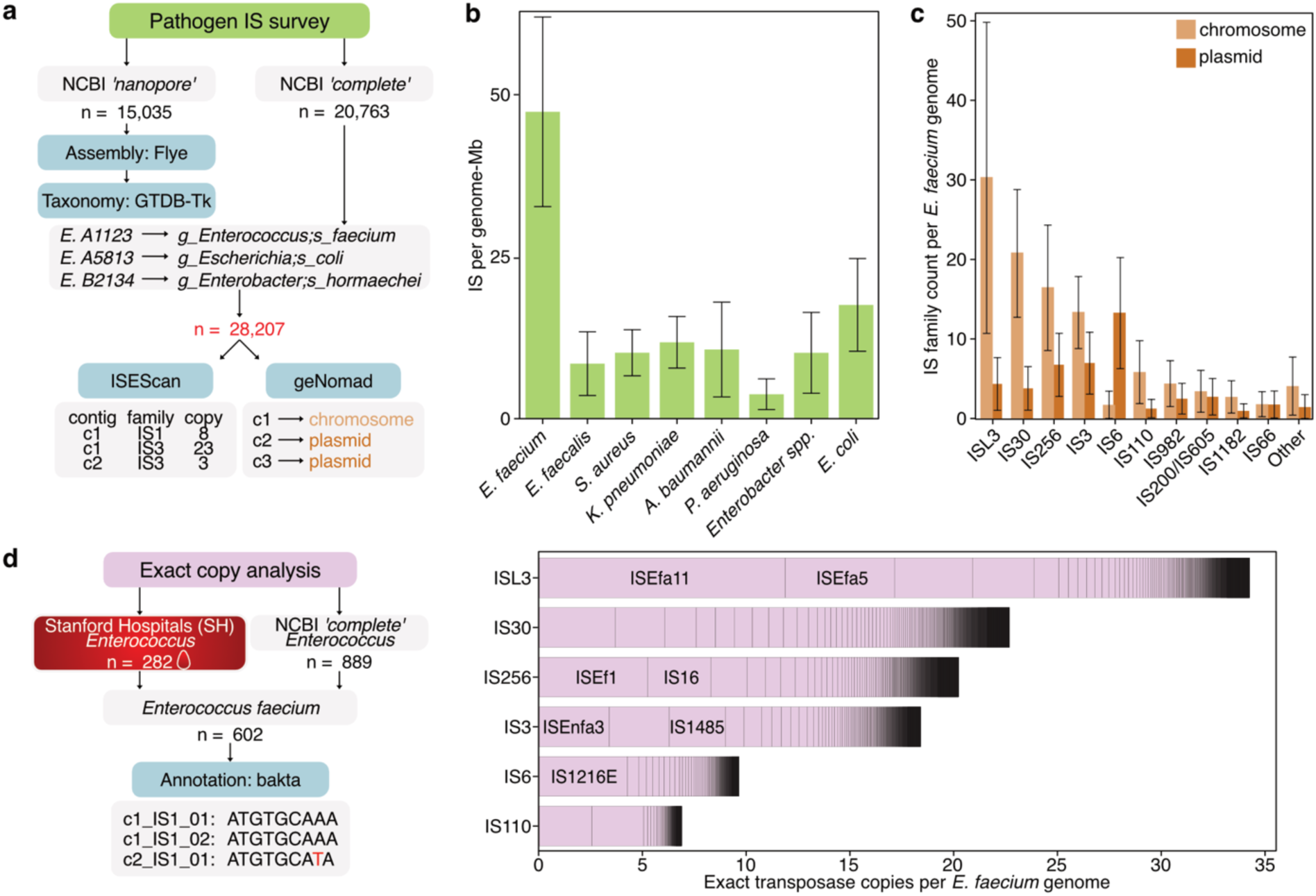
Survey of transposable elements from complete ESKAPE pathogen genomes reveals ISL3 family insertion sequences in *Enterococcus faecium* as the most abundant. **a,** Workflow schematic for counting transposable elements from publicly available ESKAPE pathogen isolate data. **b**, Among ESKAPE pathogen genomes meeting quality control criteria, mean (± s.d) total insertion sequence copy per genome normalized by genome assembly length are visualized. **c**, Among *E. faecium* genomes, mean (± s.d.) copy per IS family per genome with counts assigned to chromosome or plasmid elements as defined by geNomad. **d**, From 282 Stanford Hospitals long-read sequenced (this study) and 889 NCBI ‘complete’ *Enterococcus* genomes, 602 high quality *E. faecium* genomes were used to count exact IS transposase copies per genome at the nucleotide level. Each exact IS transposase nucleotide sequence is represented by a segment (outlined in black) and is stacked within its IS family. The segment length represents the mean number of copies of that exact nucleotide sequence per *E. faecium* genome.

Among the ESKAPE pathogens, *E. faecium* displayed the highest IS density measured as the number of elements normalized by genome length (mean: µ=47.4 copies per Mb, s.d: σ=14.9, Fig. 1B). Within *E. faecium*, ISL3 was the most abundant family (mean: µ=35.2 copies per genome, s.d: σ=21.0), followed by IS30 (24.6µ, 8.5σ), IS256 (21.7µ, 9.7σ), IS3 (19.4µ, 6.6σ), IS6 (13.2µ, 7.2σ), and IS110 (7.1µ, 4.3σ) (Fig. 1C). ISL3 elements were particularly enriched on chromosomal contigs. Consistent with IS6’s presence in the Tn1546 composite transposon conferring clinical vancomycin resistance^34^, IS6 was predominantly plasmid-borne. Intriguingly, while previous reports have indicated the importance of IS elements in the emergence of both healthcare-associated *E. faecium* and *E. faecalis*, we find that IS elements–including ISL3–are much less abundant in *E. faecalis*^16,28^ (Fig. 1B, Extended Data Fig. 1-2).

Having identified *E. faecium* as the ESKAPE pathogen with the highest IS density, we further investigated whether these IS elements exhibited signatures of recent activity. We reasoned that recently active replicative IS would appear more often as identical copies, reflecting limited time for sequence divergence^26^. To mitigate concerns about per-base sequencing accuracy, especially from earlier ONT flow-cell chemistries, here we counted exact transposase copies with available ‘complete’ genomes supplemented with 282 *Enterococcus* isolate genomes that we collected and sequenced from Stanford Hospitals (SH) (Fig. 1D). Using bakta^35^ to annotate transposase open reading frames, we found that the ISL3 transposase nucleotide sequences are more conserved than other abundant IS family transposases. Three sequences in particular, corresponding to ISEfa11, ISEfa5 and an unnamed transposase, account for more than half of the ISL3 transposase copies per genome. Notably, IS1251—a previously described ISL3 element known for its integration in the vancomycin resistance conferring *vanA* operon—is less abundant.

Taken together, these results provide a population-scale perspective on IS distributions in prominent healthcare-associated human pathogens (Extended Data Fig. 1-2). *E. faecium* is distinguished among the ESKAPE taxa by the greatest IS burden and a striking enrichment of ISL3 elements on the chromosome. The high proportion of identical ISL3 transposase copies further suggests recent replicative activity, and raises the possibility that ISL3 transposition may actively contribute to *E. faecium*’s ongoing adaptation in clinical settings.

### ISL3 elements are abundant in pathogenic Enterococci, Streptococci and Staphylococci

Prior work proposed that mobile genetic elements—including replicative IS—drove the emergence of a healthcare-associated subclade of *E. faecium* (clade A) from its commensal ancestor, *Enterococcus lactis* (formerly clade B *E. faecium*, until 2022)^17,29,36^. Clade A further separates into hospital-(A1) and livestock-associated (A2) subclades, and while one study noted enriched replicative IS3, IS110, and IS256 in A1, it did not mention ISL3. To examine the phylogenetic distribution of ISL3 among *E. faecium* clades, we utilized 805 *E. faecium* genomes from the NCBI complete, NCBI ONT and SH collections, hereafter referred to as the Complete Genome collection. Each genome was assigned to clades A1, A2, or *E. lactis* based on phylogenetic distance to reference genomes described by Lebreton *et al*^36^. We found ISL3 to be markedly enriched in the healthcare-associated A1 clade (median: 44 copies per genome) compared to A2 (2 copies) and *E. lactis* (1 copy) (Fig. 2A). This result is consistent with previous reports of relative IS enrichment in clade A1 and specifically links ISL3 to healthcare-associated *E. faecium*^17^. Notably, A1 genomes had an ISL3 interquartile range between 16 and 60 elements and a single genome contained up to 86 ISL3 elements, underscoring the considerable variability in ISL3 density even among closely related genomes, which supports the model that there is ongoing activity of ISL3 within healthcare-associated *E. faecium* (Fig. 2A).

**Figure 2.**
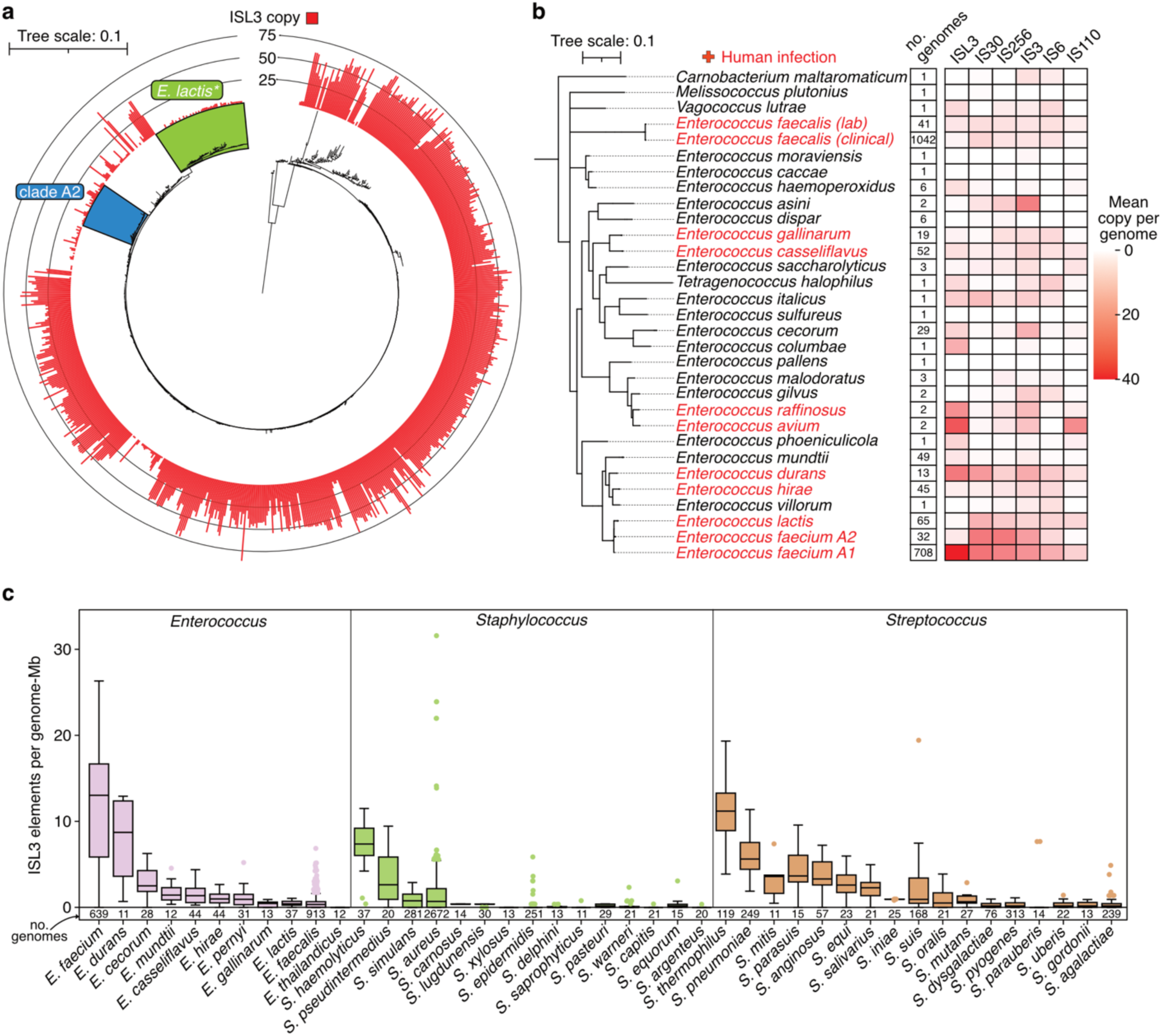
ISL3 elements are enriched in clinical *E. faecium* and abundant in other Gram-positive pathogens. **a**, ISL3 elements per genome among 805 *E. faecium* (including *E. lactis*) genomes arrayed by subclade. Genomes were assigned to clade A1, clade A2 (green), or *E. lactis* (blue, * formerly clade B) by nearest distance as measured by GTDB-Tk to the representative A1, A2 or *E. lactis* genome from Lebreton *et al*. Note that the close relationship and probable genetic exchanges between clades A1 and A2 can complicate their delineation, so some A1 or A2 assignments should be considered tentative estimates based on nearest phylogenetic distance. **b**, GTDB-Tk phylogenetic tree of *Enterococcus* genomes from the Complete Genome collection collapsed by nearest distance to the list of representative *Enterococcus* genomes from Lebreton *et al*. Heatmap values are the mean IS copy per genome among all genomes assigned to a representative. *Enterococcus* species are defined as infectious and colored red if they were positively identified to cause at least one bloodstream infection per year in the 10 year Australian *Enterococcus* Surveillance Outcome Program (AESOP). **c**, Distribution of ISL3 elements per genome-Mb across surveyed genomes for each species, separated by genus. Note, this does not include the SH or representative *Enterococcus* included in panels a and b. Boxplot boxes denote the interquartile range (IQR), thick black lines indicate the median, whiskers extend to 1.5-fold IQR, and individual values outside of 1.5-fold IQR are explicitly shown.

Non-*faecium*, non-*faecalis* (NFF) Enterococci capable of human infection are of growing clinical concern^31,37^. Intrigued by the correlation between human infection and ISL3 abundance in *E. faecium* clades, we wondered if ISL3 elements might also be abundant in NFF Enterococci. To test this, we extended our earlier approach to all 2,134 *Enterococcus* from the Complete Genome collection. These included the 282 SH isolate genomes we generated in this study, of which 8 are NFF Enterococci. We calculated the mean IS counts for the six most abundant IS families in *E. faecium* (Fig. 1C) for sets of genomes taxonomically assigned to each of 32 representative *Enterococcus* and outgroup taxa^36^ (Fig. 2B). As expected, ISL3 in clade A1 *E. faecium* was the most abundant IS element in any taxon (Fig. 2B), but we also found ISL3 elements abundant in several other Enterococci, namely *E. avium*, *E. raffinosus* and *E. durans*, all of which can cause human infection. Notably, while *E. durans* is in the same branch of *Enterococcus* as *E. faecium*, *E. avium* and *E. raffinosus* are more distantly related^38^ (Fig. 2B), supporting a model where ISL3 abundance is not strictly determined by taxonomy.

Because of ISL3’s intriguing abundance in additional infection-causing *Enterococcus* species beyond *E. faecium*, we next investigated its distribution in other clinically relevant Gram-positive bacteria. Enterococci belong to the order Lactobacillales, which also includes *Lactobacillus delbrueckii*—the earliest organism reported to harbor ISL3^39^—and *Streptococcus*, a genus that contains major human pathogens. Moreover, *Staphylococcus*, while more distantly related (within the same taxonomic class – Bacilli), also comprises numerous pathogens of clinical concern. We examined 18 *Streptococcus* and 14 *Staphylococcus* species with at least 10 available genomes in the Complete Genome collection and found that, although ISL3 was less abundant than in *E. faecium*, it remained abundant in several pathogenic *Streptococcus* and *Staphylococcus* species (Fig. 2C). Several pathogens, including *Staphylococcus aureus, Staphylococcus haemolyticus*, and *Streptococcus pneumoniae*, have abundant ISL3 elements (Fig. 2C). Of note, a subset of *Staphylococcus aureus* genomes have even more ISL3 elements than *E. faecium* (Fig. 2C). We also found *Streptococcus thermophilus*, a lactic acid producing bacteria used in commercial dairy production, as the species with the highest mean ISL3 abundance except for *E. faecium* (Fig. 2C). While ISL3 elements were generally not prevalent in the Gram-negative organisms we surveyed, they do appear in high copy in *Bordetella holmesii* (>80 per genome; Extended Data Fig. 1, Supplementary Data 1).

Together, these results support a connection between ISL3 abundance and pathogenic potential across multiple Gram-positive cocci species. As ISL3 abundance is not taxonomically restricted, this raises the hypothesis that independent acquisition and selection of ISL3 elements within a clinical context has occurred among Gram-positive pathogens.

### ISL3 elements proliferated in E. faecium over the last several decades

We were initially surprised by our observations of ISL3’s prominence in *E. faecium*: earlier reports of IS elements in the emergence of the hospital clade noted the enrichment of IS256, IS30, and IS3 elements, but not ISL3^17,29^. However, this was not an oversight, as ISL3 is not abundant in the clade A1 reference genome, which was isolated in 1998^40^. Therefore, we postulated that ISL3’s high copy (Fig. 1C) and sequence conservation (Fig. 1D) may reflect a relatively recent expansion of these elements.

Although long-read genomes allow the most accurate IS accounting, few contiguous genomes from samples collected before 2010 are available, so we developed a workflow for estimating IS copy number from short-read isolate genomic data based on read coverage (Fig. 3A). Using a collection of *E. faecium* and *E. faecalis* isolate samples with paired short- and long-read data which we generated, we first confirmed our workflow reasonably approximates genomic IS count, finding our heuristic overestimates total IS elements in short-read than long-read assemblies, by a mean 14.2% and 5.5% in *E. faecium* and *E. faecalis*, respectively (Fig. 3B). Next, we generated a temporally balanced dataset of *E. faecium* genomes sampled from before 1990 through 2024 (Fig. 3A). While IS256, IS30, and IS3 were abundant in the earliest genomes, they remained relatively stable over time, suggesting they may not be as active in the ongoing evolution of clinical *E. faecium* (Fig. 3C). By contrast, we found ISL3 elements increased from ∼10 mean copies per genome in 1990 to ∼50 per genome in 2024 (Fig. 3C). In addition, IS200/IS605 increased over the sampling window from ∼2 to ∼12 copies per genome. IS6 elements were also dynamic, which is likely due to plasmid copy number changes rather than replication of the element itself, given that IS6 elements are largely carried on plasmids (Fig. 1C), and that our short-read IS count heuristic is coverage-based. In contrast, ISL3 elements increased due to extensive replication on the chromosome (Fig. 1C). These findings demonstrate ISL3 has dramatically expanded in healthcare-associated *E. faecium* populations over the last several decades.

**Figure 3.**
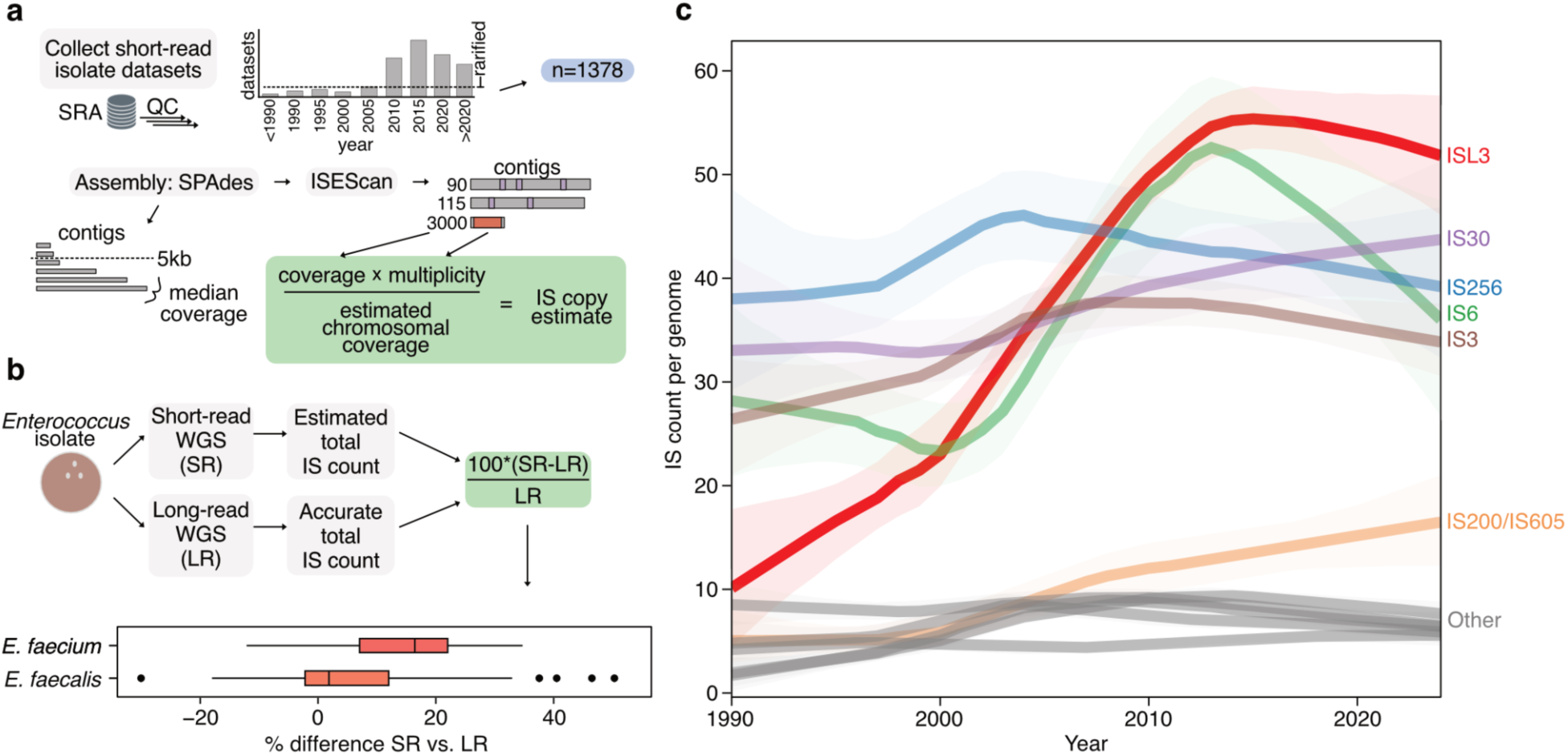
IS abundance in healthcare-associated *E. faecium* over time demonstrates recent ISL3 element proliferation. **a**, An approach to estimate transposable elements in short-read *E. faecium* data based on assembly coverage information. 39,743 available *E. faecium* isolate datasets were collected from SRA. Datasets were filtered for sufficient coverage and access to geographic location and time of collection. Then, datasets were rarified after 2005 to obtain a balanced set of 1,378 genomes assembled with SPAdes and annotated with ISEScan. For each dataset, IS count per family was estimated by normalizing the coverage and IS multiplicity for a given contig by the estimated chromosomal coverage. **b**, Isolate genomes (38 *E. faecium*, 81 *E. faecalis*) were sequenced with both short-(SR) and long-reads (LR) and assembled with SPAdes or Flye, respectively. The percent difference between the total IS counts from the resulting assemblies is shown as a boxplot with the median, 25th and 75th percentiles, and individual values outside of 1.5 times the interquartile range visualized. **c**, Using the approach outlined, the mean insertion sequence copy per genome binned by sample year from 1990 until 2024 (all data before 1990 is binned with 1990) is shown as a distinct LOWESS trendline with 95% confidence intervals in the same color at reduced saturation.

### Extensive structural variation in hospital E. faecium is largely mediated by ISL3 replication

Whether ISL3 continues to be active in present-day *E. faecium* is unclear. Case studies in longitudinally collected clinical isolates of *E. faecium* have measured IS element transposition activity within a single patient^41,42^. However, the broader landscape of IS-driven genome plasticity within a present-day endemic hospital lineage infecting many patients remains undefined. To address this gap and to investigate how ISL3 might underpin the success of *E. faecium* in contemporary clinical settings, we sought to determine 1) the extent of short-term structural variation within a single hospital lineage of *E. faecium* and 2) whether ISL3 activity contributes to any observed variation.

To accurately detect ongoing genomic structural variation, highly-contiguous and minimally-divergent genomes are necessary. Therefore, we collected every bloodstream isolate of *E. faecium* (n=109) from Stanford Hospitals (SH) in 2021 and 2023, performed long-read sequencing, assembled complete or near-complete genomes, and identified clusters of highly-related genomes using PopPUNK^43^ (Fig. 4A). Consistent with epidemiological studies from other institutions^44–46^, a single dominant lineage of *E. faecium* was present. At Stanford Hospitals, sequence type (ST) 117 accounted for 66/109 isolates (Fig. 4B). This ST117 cluster represents an ideal population to investigate recent IS activity within a hospital setting.

**Figure 4.**
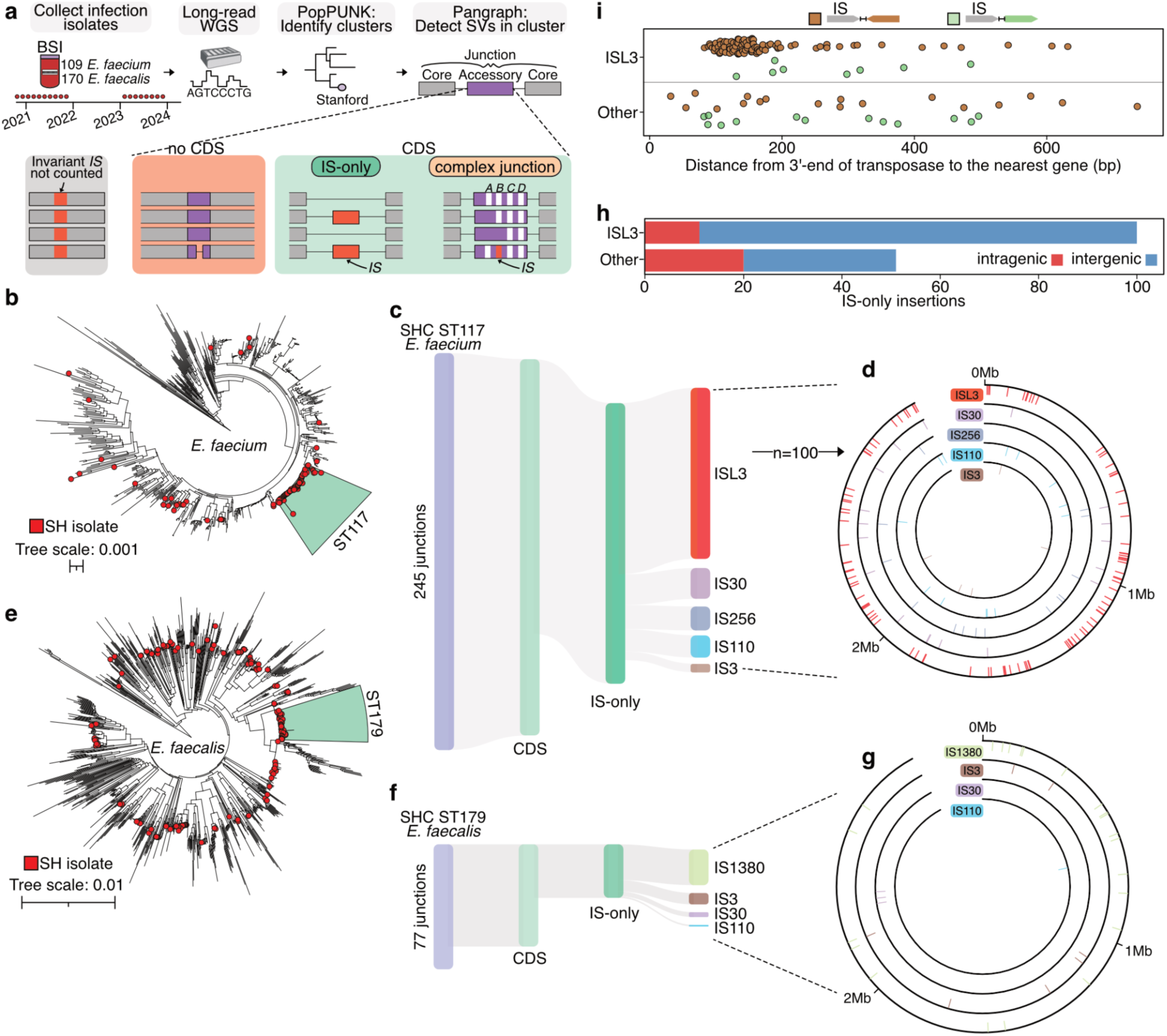
ISL3 mediates extensive recent structural variation within a hospital-endemic *E. faecium* lineage. **a**, Analysis of recent IS activity among a closely related hospital lineage using Pangraph. **b**, PopPUNK clustering and phylogeny of *E. faecium* isolates from Stanford Hospitals (this study; red tips) among publicly available genomes. An *E. faecium* cluster of ST117 genomes is highlighted (n=66). **c**, IS variants (164/245 total) detected in SH ST117 *E. faecium*. **d**, Circos plot showing the genomic location of each IS variant from panel c. Rings are ordered and elements colored the same as panel c with ISL3 variants in the outermost ring and IS3 in the innermost. **e**, PopPUNK clustering and phylogeny of *E. faecalis* isolates from the Stanford Hospital collection (this study; red tips) among publicly available genomes. An *E. faecalis* cluster of ST179 genomes is highlighted (n=44). **f**, IS variants (33/77 total) detected in SH ST179 *E. faecalis*. **g**, Circos plot showing the genomic location of each IS variant from panel f. Rings are ordered and elements colored the same as panel f, with IS1380 elements in the outermost ring, and IS110 elements in the innermost. **h**, Among simple insertion variants in SHC ST117 *E. faecium*, the number of inter- vs. intragenic insertions for ISL3 and the other IS elements named in panel c is displayed. **i**, For both of these categories, the distance from the 3’ end of the transposase coding sequence to the next gene is displayed. Each dot is colored by whether the IS is oriented in the same (green) or opposite (brown) direction as the downstream gene.

As a control, we employed the same approach for *E. faecalis* (n=170), a species recognized for previous IS expansion^16,28^, with similar genome size and causing a similar number of infections over the same time period at the same hospital (Fig. 4A). We identified a cluster of *E. faecalis* genomes from 44 SH isolates corresponding to ST179 (Fig. 4D). Importantly, the recombination-filtered core genome divergence of the ST179 *E. faecalis* and ST117 *E. faecium* clusters are comparable (∼.0009 and ∼.0005 SNPs per core genome-aligned Mb, respectively), indicating they have experienced similar rates of SNP accumulation (Extended Data Fig. 3). Therefore, rates of structural variation between the clusters can be directly compared, allowing us to weigh the relative importance of recent structural variation between these pathogens.

Within the *E. faecium* ST117 cluster and the *E. faecalis* ST179 cluster, we measured recent IS activity by employing a reference-free, gene-agnostic approach based on that described by Molari *et al*^47^ (Fig. 4A). This method allows us to detect structural variations (genomes with IS elements in different positions) without the biases introduced by reference-based approaches, providing a comprehensive view of genomic plasticity within this constrained lineage (rather than compared to a single reference). Within the ST117 cluster, we identified 245 structurally variable loci (Fig. 4C-D). IS-mediated insertions accounted for 67% of these junctions, with ISL3 predominating the landscape of recent activity (Fig. 4C-D). In contrast, the ST179 *E. faecalis* cluster showed markedly fewer IS-mediated structural variations (Fig. 4F-G), supporting a model where recent IS activity plays a more prominent role in the ongoing evolution of *E. faecium*. This analysis also identifies IS elements that contribute relatively little to ongoing evolution in *E. faecium*. While IS30, IS256, IS3 and IS110 elements are collectively much more numerous than ISL3 elements (Fig. 1C), they contribute more than 50% fewer recent variants (Fig. 4H).

As the genome reductive effects of IS expansion have been repeatedly observed, we next investigated whether these active ISL3 insertions might also be leading to insertional disruption. Among 100 distinct ISL3 insertions observed within ST117, only 11 disrupted open reading frames, a markedly lower proportion than for other common IS families (Fig. 4H). Instead, ISL3 elements are repeatedly inserted into intergenic regions, typically on the opposite strand of— and very close to—the nearest downstream gene (Fig. 4I). These findings highlight the striking genome plasticity taking place over a few years in a single hospital-endemic lineage of *E. faecium* driven by active ISL3 transposition.

### *In situ* identification of structural variation from single patients with long-read metagenomics

*E. faecium* is known to colonize and often dominate the gastrointestinal tract of individuals with hematological malignancies including those undergoing hematopoietic cell transplantation (HCT)^20,41^. Gut domination also often precedes bloodstream infections with this pathogen^48,49^. Consistent with a pattern of short-term adaptation, previous studies have reported SNP level changes in *E*. *faecium* during intestinal domination and the transition to bloodstream infection^3,50–52^. Given the high copy number of ISL3, evidence of its high levels of active replicative insertion, and its capacity to facilitate recombination and subsequent structural variation, we sought to measure whether replicative ISL3 activity might induce observable structural variation in *E. faecium* within the gut microbiomes of single patients over time. To investigate this hypothesis, we curated a set of 34 longitudinal stool samples from 14 patients who underwent HCT at Stanford Hospital^49,53^ and who had evidence of longitudinal *E. faecium* domination, defined here as multiple samples with >10% relative abundance (Fig. 5A, Extended Data Fig. 4). Importantly, isolation and culturing of *E. faecium* from stool introduces a severe population bottleneck, potentially introducing or selecting for rare variants. To overcome this limitation, we directly long-read-sequenced stool samples, enabling evaluation of near-native population frequencies of structural variants without the bias introduced by isolation and culture. After excluding two samples for poor DNA extraction and one sample for low sequencing depth, the remaining 31 samples had a median read quality of Q23.1 and a median read N50 of ∼9.18 Kb (Extended Data Fig. 4, Fig. 5A). After assembly and taxonomic profiling, we identified a single *E. faecium* chromosomal contig for 28/31 samples (Extended Data Fig. 5B-C).

**Figure 5.**
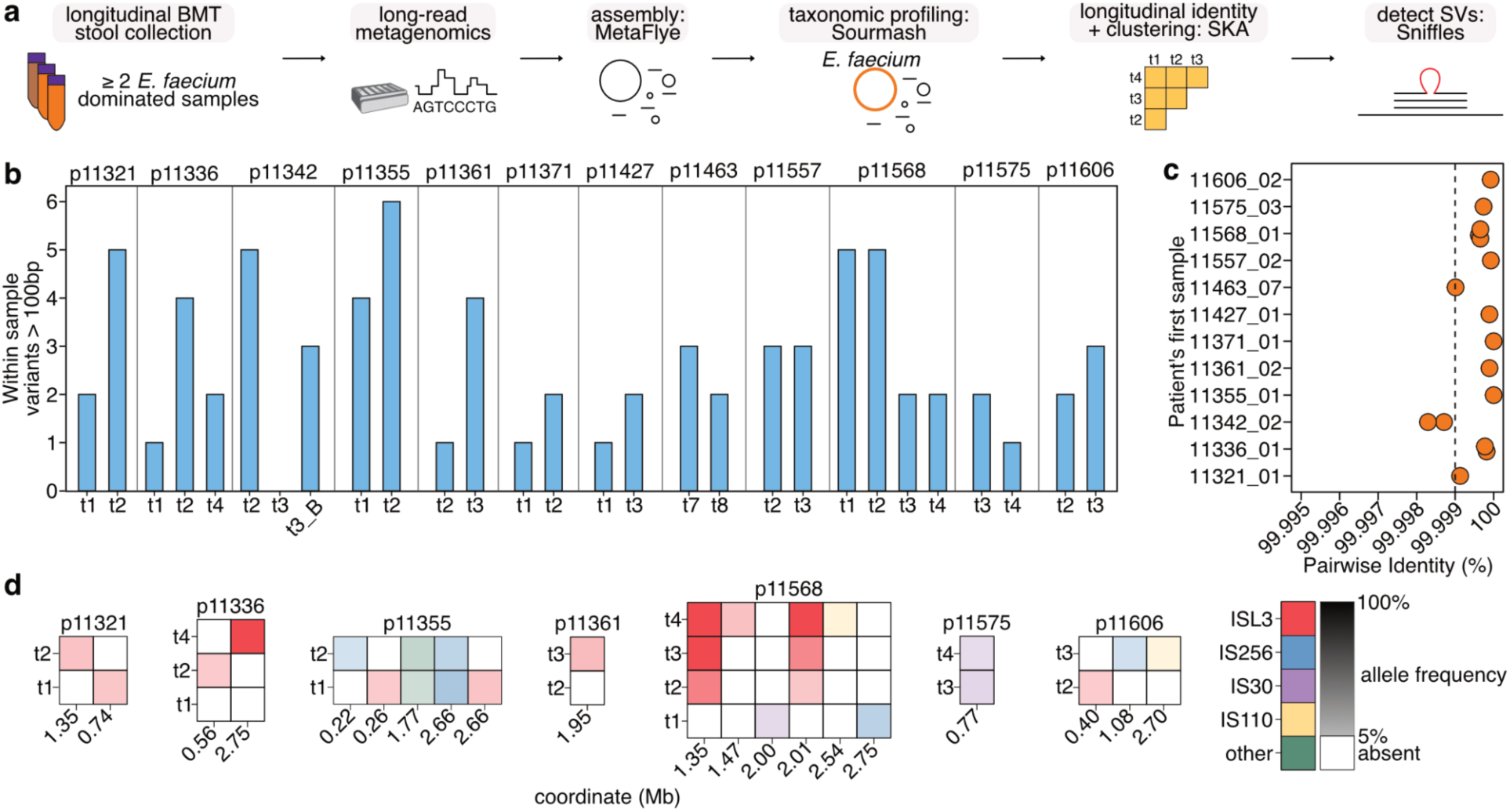
Longitudinal long-read metagenomics of patient stool samples identifies structural variants in otherwise isogenic strains of *E. faecium*. **a,** A workflow for performing longitudinal long-read metagenomic sequencing to identify structural variation of *E. faecium* against an isogenic background. **b**, Within-sample heterogeneity of 28 *E. faecium* chromosomes (Extended Data Fig. 4, 5). The number of variants per sample longer than 100bp predicted by Sniffles are shown. Samples are grouped by the anonymized patient ID^53^ (pXXXXX) and labeled by the time point for that patient. **c**, For the 12 patients with multiple samples having a complete *E. faecium* contig (Extended Data Fig. 5), the pairwise identity of *E. faecium* contigs from the same patient as measured by SKA, relative to the first dominated sample is shown. Samples are labeled in the format pXXXX_timepoint. The identity threshold of 99.999% or 10 SNPs/Mb is indicated with a dashed line. **d**, Among the 11 patients reaching the identity threshold, IS variants predicted by Sniffles are shown per patient and arrayed by their patient-specific genomic coordinate (Mb). The color represents the IS family and the saturation represents the allele frequency.

Within a single gut-resident *E. faecium* population, variants may be observed at intermediate frequencies, reflecting alleles that have not fixed in the population; these alleles may change in frequency or fix over time, reflecting selection and/or drift. Such population dynamics may not be visible in genome assemblies, which collapse population heterogeneity, yet this information may be latent in sequencing data with sufficient depth. To detect these variants, we used Sniffles2^54^, a tool for calling structural variants, including IS variants, from long-read sequencing data by comparing the raw reads to a reference genome. By comparing the reads from each sample to their own *de novo* assembly, we detected evidence of within sample *E. faecium* structural heterogeneity in 27/28 stool samples (Fig. 5B).

Next, we looked for IS variants that changed in frequency over time in individual *E. faecium* patient populations. To first confirm strains remained identical at the SNP level over time, we measured pairwise genetic identity between *E. faecium* chromosomes with SKA^55^ (Fig. 5A). Among 12 patients with longitudinal samples (Extended Data Fig. 4, 2 patients became singletons), and using a threshold of 10 SNPs/Mb, we confirmed 11 patients had ‘isogenic’ strains over time, with p11342 as the single exception (Fig. 5C). Considering all pairwise distances, we also observed two clusters containing *E. faecium* from multiple patients indicative of strain sharing (Extended Data Fig. 5D). One cluster corresponded to the same ST117 lineage (Fig. 4B) present at Stanford Hospitals from 2021-2023 (Cluster 1, Extended Data Fig. 5D). Given the earliest stool sample in Cluster 1 (p11321_01) was collected in 2015, this suggests the Stanford ST117 *E. faecium* lineage has been present in the hospital environment for at least 8 years, to our knowledge the longest duration example for *E. faecium*. In Cluster 2, all pairwise comparisons between p11371 and p11427 samples had fewer than 8 SNPs (Extended Data Fig. 5D). Metagenomic samples from these patients were previously analyzed for clinical transmission with short- and linked-read technologies, yet incomplete assemblies prevented their confident identification^53^.

For each patient with confirmed strain identity across all timepoints, we aligned the raw sequencing reads from each sample against the metagenomic assembly from that patient’s earliest time point. We then filtered the resulting variants that aligned to the *E. faecium* chromosome for predicted IS element insertions. IS variants were dynamic in 7/11 longitudinal cases (Fig. 5D), including one patient, p11568, in which we observed two ISL3 insertions monotonically increasing in frequency over time across p11568’s four samples.

This approach identifies IS-mediated structural variation in natural populations of clinical pathogens, capturing population heterogeneity within samples that may otherwise be confounded by isolation and lost during assembly. Furthermore, considering longitudinal dynamics of IS elements, especially in the context of clinical metadata, can reveal variants being acted upon by selection in the clinical environment.

### A recurrently observed ISL3 insertion near *folT* activates folate scavenging, conferring a fitness benefit

Intrigued by the ISL3 mediated variants reaching high allele frequency over time in p11568, an individual who had undergone allogeneic hematopoietic cell transplantation for treatment of acute myeloid leukemia (Fig. 5D), we further examined this case to investigate the effects of recent ISL3 structural variants on the *E. faecium* genome. Four stool samples with *E. faecium* domination were collected on days -2, 12, 19 and 26 (time point 1-4; t1-t4) relative to transplant (day 0). Examining these insertions closely, we find that one ISL3 variant inserted 42 bp upstream and created a 10 bp repeat in the same orientation as a folate transporter (*folT*) (Fig. 6A-C). The second ISL3 variant disrupted *agrC* within the accessory gene regulator operon (*agr*) (Extended Data Fig. 6A-C). Neither variant was detectable in t1 but both increased in frequency to fixation by t4 (Fig. 6A, Extended Data Fig. 7). We hypothesized the *folT* adjacent variant has a regulatory effect on the expression on *folT*. To investigate this hypothesis, we isolated *E. faecium* from t1 and t4 stool. We found the strains to be nearly identical (no SNVs or indels), with the only detectable differences being the difference in the position of two ISL3 elements, as described above, and a presumed phage element that is present in t4 but not t1. With the insertional genotypes confirmed, we performed RNA-seq (Fig. 6B, Extended Data Fig. 6, 7). Strikingly, we found that *folT* is the most upregulated gene in the genome (Fig. 6B, Extended Data Fig. 7), while the disrupted *agr* operon is silenced in t4 (Extended Data Fig. 6, 7). Upon closer examination of the ISL3 insertion upstream of the *folT* locus, we found that the locus retains its native -10 promoter sequence (Pribnow box) (Fig. 6C). However, the insertion replaces the native -35 sequence (5’-ATGACA-3’) with a ‘perfect’ -35 sequence donated by the ISL3 element (5’-TTGACA-3’) (Fig. 6C). These findings demonstrate that the ISL3 element carries a 3’-outwardly directed -35 promoter sequence that can drive expression when positioned correctly upstream of a Pribnow box and transcriptional start site.

**Figure 6.**
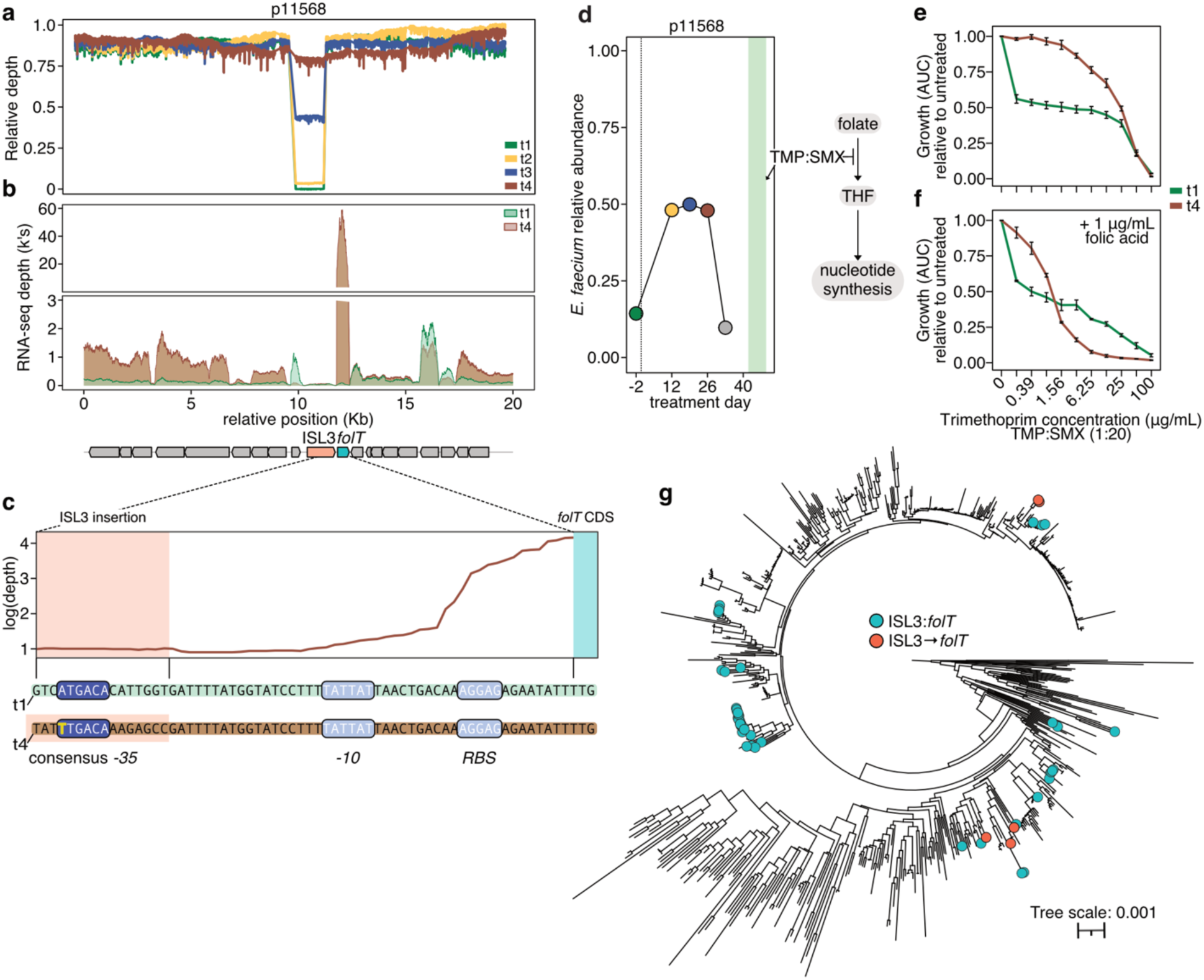
A high frequency ISL3 structural variant emerges, accompanies *E. faecium* gut domination and drives folate scavenging. **a**, In the neighborhood of the ISL3 variant fixing upstream of *folT*, the relative coverage of long-reads aligned to the metagenomic assembled *E. faecium* chromosome of p11568 t4, normalized to the average chromosomal coverage per sample. The coordinates are relative to a 20 Kb neighborhood centered on the ISL3 variant. **b**, From *E. faecium* isolated from p11568 t1 and t4 stool samples, RNA-seq read coverage (log-scale) across the same locus as in panel a. **c**, The sequence context of the ISL3 variant hybrid promoter is shown in detail. The mean depth of t4 isolate RNA-seq reads (n=4) aligned to the t4 isolate assembly in log-scale from near the 3’-end of the ISL3 insertion (red background) to the start of the *folT* CDS (cyan background) is shown. The aligned assembly sequence from the t1 and t4 isolates are shown below, with putative *-10* and ribosome binding sites (*RBS,* light blue) and *-35* promoter elements (dark blue) highlighted. **d**, Relative abundance of *E. faecium* in t1-t4 stool samples as estimated from previous short and linked-read metagenomics^53^ for p11568. Samples from day -2, 12, 19 and 26 relative to transplant correspond to t1-t4. Transplant day 0 is indicated with a dashed line. p11568 received trimethoprim-sulfamethoxazole (TMP:SMX) between day 42 and 49 after transplant. (right) A simplified schematic showing where TMP:SMX acts within the folate activation pathway. **e**, Relative to untreated, the growth (area under the curve, AUC) of the t1 and t4 isolates in increasing concentrations of TMP:SMX (concentration of TMP is labeled, TMP:SMX was prepared in a ratio 1:20). **f**, Same as panel e, with all conditions supplemented with 1 µg/mL folic acid. **g**, The incidence of ISL3 insertions co-oriented, and upstream of *folT* (orange) and adjacent to *folT,* but in different orientations (blue), among a phylogeny of 687 *E. faecium* genomes with *folT* as measured by PopPUNK.

Healthcare-associated *E. faecium* is clinically resistant to antifolate antibiotics, such as trimethoprim, via the uptake of exogenous folates^56^. We wondered if the increased expression of *folT,* which encodes the substrate-specific component of an ECF folate transport complex^57^, might have been selected for by treatment with antifolate medications. However, review of p11568’s clinical metadata revealed no evidence of exposure to either methotrexate (an anticancer agent inhibiting folate metabolism) or trimethoprim-sulfamethoxazole (TMP:SMX) during the time frame of our sampling (Fig. 6D). Therefore, we reasoned that increased *folT* expression had some other adaptive benefit, such as enabling better competitive growth of the folate auxotrophic *E. faecium* in the setting of limited folate in the gut. To test the effect of increased *folT* expression on the competitive fitness of p11568 t1 and t4 isolates, we measured their growth *in vitro* during treatment with TMP:SMX in order to determine whether exogenous folate in the media could be differentially utilized to rescue the toxic effects of antifolate drugs (Fig. 6E-F, Extended Data Fig. 8). Indeed, we found that the t4 isolate had a marked growth advantage over t1 at all tested drug concentrations (Fig. 6E) and this difference was entirely abrogated when the media was supplemented with the synthetic folate folinic acid, which bypasses antifolate toxicity (Extended Data Fig. 8). Conversely, we observed that supplementation with folic acid produced increased drug susceptibility in both isolates and particularly in t4, which is likely due to more rapid accumulation of non-productive folate intermediates^58^ in this isolate (Fig. 6F). Together, these findings suggest increased expression of *folT* driven by ISL3 activity enhances p11568 t4 *E. faecium*’s ability to scavenge folate metabolites from the environment, thereby conferring a fitness advantage over p11568 t1 under folate limiting conditions.

While it is possible that a variant, such as the *folT*-ISL3 variant observed here, may reach high frequency in an individual strain due to genetic drift, observing it repeatedly in multiple distinct strains and/or with multiple distinct genotypes provides compelling evidence for selection. As *folT* expression appeared to have clinically relevant effects in p11568, we reasoned that there may be other examples of IS insertion near *folT*. Returning to our collection of global healthcare-associated *E. faecium* (Fig. 4B), we investigated the neighborhood of the *folT* locus. In 50 of the 687 genomes with a *folT* locus, ISL3 was present directly adjacent to *folT,* which might point to the *folT* locus being an insertional hotspot (Fig. 6G). In five of these genomes, as in p11568 t4, an ISL3 element was positioned upstream of, and co-oriented with, *folT*. These ISL3→*folT* variants were found in bloodstream infection or HCT patient gut isolates in the Netherlands, Germany, and South Korea. Occurring across multiple phylogenetic lineages and involving different ISL3 copies, they likely represent independent evolutionary events. Given their global spread, these variants implicate folate scavenging as an important function under selection in critical care patients.

## Discussion

Healthcare-associated pathogens that reside in the gut microbiome are under incredible environmental pressure to evolve and adapt, both to antimicrobial agents and to metabolic challenges that arise from collapse of cross-feeding interactions. While SNPs and small indels most certainly contribute to this adaptation, phenotypic variations caused by selfish genetic elements are likely major and under-appreciated contributors to these types of adaptation. Here, we report the breadth of the recent replicative ISL3 expansion in the healthcare-associated pathogen, *Enterococcus faecium,* and implicate its importance in several other major Gram-positive pathogens. Furthermore, we demonstrate the ongoing, extensive structural variation mediated by ISL3 elements in a deeply sampled contemporary *E. faecium* lineage endemic to a hospital. Focusing on hospitalized individuals undergoing HCT, we use long-read metagenomics to identify physiologically relevant structural variation and population heterogeneity within *E. faecium* dominating single-patient guts. In one patient’s gut-dominant *E. faecium* strain, two ISL3 variants rise to high frequency. One variant generates a hybrid promoter, driving the expression of *folT*, which we find repeatedly in clinical *E. faecium* strains, likely contributing to improved competitive fitness through upregulation of a transporter for a scarce and essential micronutrient.

Previous efforts to survey IS content and diversity in prokaryotic genomes relied on available complete genomes distributed broadly across the bacterial kingdom numbering in the tens to hundreds^24–26^. Here, leveraging the availability of tens of thousands of complete genomes, we investigated IS distributions in healthcare-associated pathogen taxa. We find *E. faecium* is particularly enriched for IS elements among healthcare-associated pathogens (Fig. 1B). While an enrichment of mobile genetic elements has been noted for several pathogens, including *E. faecium*, this survey places the relevance of IS elements within pathogen genomes into context (Fig. 1B, Extended Data Fig. 1). While *E. faecium* has the most IS elements, subsets of other pathogens are rich with IS elements, including *A. baumanii* and *S. aureus,* raising the possibility these organisms are poised to undergo similar evolutionary processes (Extended Data Fig. 1). Within *E. faecium* we found replicative ISL3 elements are the most abundant (Fig. 1C). Given ISL3 elements were first described in *Lactobacillus*^39^, albeit at a much lower copy number, it is unsurprising that we observed ISL3 elements in *Enterococcus*, another member of Lactobacillales. This also might explain the identification of ISL3 in *Streptococcus pneumoniae* and *Staphylococcus* species including *S. aureus,* as these organisms share the same class (Bacillli). Interestingly, it appears that ISL3 is not strictly taxonomy bound; for example, ISL3 elements have been reported in Gram-negative organisms^20^ and we observe them to be abundant in *Bordetella holmesii* within our dataset, suggesting they may have a much broader taxonomic scope (Extended Data Fig. 2). This suggests that ISL3 is either an ancient element that has diverged into different subclasses over time, or that it can be actively transmitted through elements such as a plasmid, which has been observed for *K. pneumoniae*^59^. While ISL3 is highly abundant in nearly all *E. faecium* clade A1 strains, these elements are not abundant across all species of *Enterococcus, Streptococcus* and *Staphylococcus.* This suggests features that activate, drive or suppress IS activity are differentially present across these organisms, with clade A1 *E. faecium* either harboring an activator or driver or lacking a suppressor of these elements. Such a driver or suppressor could be a gene; however, another potential driver of ISL3 activity might be the clinical environment. While we found a number of species in these taxa with few or no ISL3 elements, those species with high ISL3 abundance are often those capable of causing human infection, including *Streptococcus pneumoniae, Staphylococcus aureus* and *haemolyticus,* and *Enterococcus durans, avium* and *raffinosus*.

In contrast to other, well-described and abundant IS elements in healthcare-associated *E. faecium*, such as IS256^29^, we find that ISL3’s copy number has increased substantially in the past 30 years. This could be explained by an initial burst of copy number followed by ‘silencing’ of the IS element and maintenance of the elements in a fixed position^26^. Using a reference-free, Pangraph-based approach^47^ to analyze a collection of long-read sequenced isolates we generated from a single hospital over three years, we find that this is not the case; rather, we find that ISL3 positions within *E. faecium* genomes are highly variant, and much more so than other abundant IS elements. While others have demonstrated, *in vitro*, that elements such as IS256 can be variant in position in sequential bacterial isolates taken from patient bloodstream infections^42^, our results suggest ISL3 is much more active than the other IS elements within *E. faecium*. Furthermore, given ISL3’s relative enrichment in intergenic vs. intragenic locations, it appears that its impact on bacterial fitness and physiology is not largely driven by insertional disruption of genes. Instead, we suspect the observed propensity for intergenic insertion may reflect its tropism to relatively AT-rich intergenic regions or the favorable selection of intergenic insertions with regulatory impacts compared to the more likely deleterious effects of intragenic disruption. Consistent with this, prior work has demonstrated examples of outwardly directed promoter elements on IS that can drive transcription of neighboring genes and adaptive fitness benefits such as antibiotic resistance^60,61^. Continued surveillance of ISL3 and its copy number, in *E. faecium* and other organisms that harbor it, will likely inform our understanding of its ongoing rate of activity and may point toward genes that might participate in regulating its activity.

Given the curious dominance, persistence, and recent copy number increase of ISL3, it is interesting to consider the underlying reasons for its relative success. Selfish genetic elements that occasionally provide a regulatory function that can be ‘toggled’ on and off — such as the *folT* promoter alteration we observed — may evade purifying selection and persist under strong clinical pressures. By donating a ‘perfect’ -35 promoter sequence, this element can take genes that are less highly expressed due to a native ‘imperfect’ -35 promoter sequence and can dramatically upregulate expression of genes downstream of this locus. We identified multiple genomes with *folT*-proximal ISL3 insertions that occurred in the same configuration as we observed in p11568 t4. As *folT* is a hotspot for ISL3 insertion generally, future work is needed to understand the potential regulatory impacts of other configurations. This type of benefit may enable both *E. faecium* and the ISL3 element to evade negative selection. This can then lead to accumulation of these elements throughout the genome, especially in situations where competition is scarce^62^. Because identical elements in multiple sites within a genome are a nidus for recombination, a consequence of ISL3 expansion might be deletion of regions of non-essential regions of the genome. This type of genome reduction has been described in the past — having occurred through the sequential expansion of an IS element followed by reductive evolution — in *Bordetella pertussis*^10^, *Yersinia pestis*^9^ and in the evolution of *E. coli* into *Shigella* lineages^14^. In each of these cases, genome reduction contributed to a narrower host range, and a more pathogenic lineage with significant impacts on human society. For example, *Shigella*, unlike *E. coli*, is effectively an obligate human pathogen and has no known reservoirs outside of humans. The notion that *E. faecium* may be following a similar path toward hospital specialization is supported by the observation that clade A1 strains dominate the gastrointestinal tract of critically ill patients but have limited prevalence in healthy gut microbiomes^16,48^. It is intriguing to note that direct evaluation of clade A1 *E. faecium*’s major reservoir, the gut microbiomes of hospitalized patients, provides direct long-read metagenomic evidence of multiple (up to 6) ‘IS-variant strains’ in a single gut microbiome. This direct evaluation of the natural population of *E. faecium,* without the bottlenecking effect of isolation and culture, enables important insight into the importance of ISL3 to *in vivo E. faecium* evolution within the clinical environment.

Our work has several limitations. First, our Complete Genome collection likely underestimates the population dynamics of transposable elements in at least three ways: 1) every sample experienced the severe bottleneck of isolation and population heterogeneity is collapsed during assembly 2) the available complete genomes are highly non-random geographically and phylogenetically, 3) the majority of available complete genomes are from the last decade and represent collections of isolates from generally resource-rich hospitals or academic centers. Our choice to focus on *E. faecium* may have been influenced by this limitation as the available complete genomes of *E. faecium*, in contrast to more broadly studied *S. aureus* or *E. coli,* are largely restricted to clinical samples from hospitalized inpatients. Indeed, for all of the ESKAPE pathogens, we observe some subset of genomes with extensive IS proliferation (Extended Data Fig. 1-2). Second, although we find extensive evidence of IS mediated structural variation in the Stanford Hospital endemic ST117 lineage, we sampled only bloodstream infection isolates. This collection, again potentially confounded by isolation, does not include samples from hospital surfaces or from patient guts, the main reservoirs of *E. faecium* in the hospital. Finally, we analyzed structural variants mediated by chromosomal transposable elements with Pangraph. Plasmids are known to both be enriched in transposable elements that can drive chromosomal integration of IS elements^59,63^ and important mediators of *E. faecium* pathogenicity^64–66^. While our approach does capture structural variants of integrated prophages, our approach does not capture plasmid rearrangements.

In conclusion, we find replicative ISL3 elements are a major driver of current and ongoing evolution in the healthcare-associated pathogen, *Enterococcus faecium*. With the rapidly increasing availability of contiguous genomes, broadening genomic surveillance across pathogens and other related species is essential for capturing the dynamics of these elements over time and across different environments. We believe these efforts may help anticipate the emergence of new pathogens and identify human behaviors involved in their spread. It remains unclear what element-intrinsic or host-extrinsic factors contribute to the success of ISL3 elements. Understanding these factors will be vital in developing strategies to limit their evolvability and manage their impact on pathogenicity. Furthermore, considering evidence of this relatively nascent ISL3 expansion through a population genetic framework, it is possible that *E. faecium* is undergoing the early phases of the same sort of IS-mediated transformation process that *Shigella*, *Bordetella* and *Yersinia* have in the past. By elucidating these mechanisms of genomic plasticity, our study offers novel insights into how bacterial pathogens leverage mobile genetic elements to swiftly respond to the intense selective pressures of clinical settings. Such understanding not only advances our knowledge of bacterial evolution but also holds promise for developing more effective strategies to combat the rising threat of healthcare-associated infections.

## Methods

### NCBI ONT collection, preprocessing, assembly, and quality control

Candidate isolate long-read sequencing datasets were identified on NCBI with the following search criteria on February 29th, 2024: ((((Enterococcus[Organism]) OR (Staphylococcus[Organism] OR (Klebsiella [Organism]) OR (Acinetobacter [Organism]) OR (Pseudomonas [Organism]) OR (Enterobacter [Organism]) OR (Escherichia [Organism]) OR (Salmonella[Organism]) OR (Streptococcus[Organism])) AND “oxford nanopore”[Platform]))). Datasets for *Shigella* and *Bordetella* were separately queried on October 28th, 2024 in the same way. Metadata for these accessions was downloaded with the SRA Run Selector interface. Datasets with less than 50 Mb of total sequencing or less than 2kb average read length were not included for further analysis. Raw reads were downloaded using SRA tools prefetch and fasterq-dump, and assembled with flye v2.9 using the `--nano-raw` method. Genome taxonomy was verified with gtdb-tk v2.4.

Assemblies were then filtered for appropriate contiguity (<=20 contigs), genome size (+/- 2 SD per genus), coverage (>=40x), and with identified species-level taxonomy. We excluded the GTDB-Tk taxonomy result for the *Shigella* datasets because the GTDB-Tk database does not include *Shigella* genomes and thus cannot classify them beyond *E. coli*. Finally, we filtered the set of genomes to only include species with >10 samples in this and the NCBI complete collection (below) combined. The final set of genomes with assembly and taxonomy information is available in Supplementary Data 1, and the custom scripts used to filter and define the final set of genomes to produce the table are available in the `data_processing` module of our GitHub repository.

### NCBI complete genomes

The NCBI command line tool ncbi-datasets-cli v16.22.1 was used to download all available NCBI genomes with the ‘complete’ assembly level from the following taxa on July 3, 2024: *Enterobacter, Staphylococcus, Klebsiella, Acinetobacter, Pseudomonas, Enterococcus, Escherichia,* and *Salmonella, Streptococcus, Shigella,* and *Bordetella*. For example, the following command was used to download *Enterococcus* assemblies: `datasets download genome taxon enterococcus --assembly-level complete --include genome,gff3 -- dehydrated --filename enterococcus.zip`. After rehydrating, duplicate assemblies were removed, preferring the RefSeq over Genbank as the source database. Assemblies were then filtered according to the same contiguity, size, coverage, and sample count thresholds as above.

### Stanford Hospitals (SH) *Enterococcus* isolate collection and sequencing

279 bloodstream infection isolates identified by biotyping with a MALDI-TOF based Bruker Biotyper as *Enterococcus faecalis* (n=170) or *Enterococcus faecium* (n=109) by the Stanford Healthcare Clinical Microbiology Laboratory between the periods 2020/10/01-2021/12/31 and 2023/01/01-2023/12/31 were collected under an IRB protocol approved by The Stanford University Research Compliance Office (Protocol #64591; Principal Investigator: Dr. Ami Bhatt). An additional 8 isolates collected in 2024 identified as atypical *Enterococcus* as above were stored as glycerol stocks. Metadata including date, site of collection, and relevant patient information were extracted. Each isolate was cultured aerobically for approximately 16 hours with shaking at 37°C in 3 mL Brain Heart Infusion (BHI) broth (Sigma-Aldrich). Then, a 1:100 subculture into 10 mL BHI broth was prepared. Subcultures were allowed to reach OD ∼1.0. The subculture was centrifuged at 3,000xg for 10 minutes at room temperature, resuspended in PBS, then centrifuged again. The PBS was poured off and the subsequent pellet was resuspended in 1 mL 1x Zymo DNA/RNA Shield (Zymo: R1100-250) and shipped to Plasmidsaurus (Oregon, USA) for ONT ‘Standard Bacterial Genome with DNA extraction’ sequencing. Upon inspection of the assemblies, one *E. faecium* isolate (EF_B_40) was mischaracterized as *E. faecalis* and one *E. faecalis* isolate (FM_B_40) was mischaracterized as *E. faecium* based on biotyping. Of the 170 *E. faecalis* isolates, 167 were confirmed as *E. faecalis*. Of the 109 *E. faecium* isolates, 107 were confirmed as *E. faecium* with standard quality control performed by Plasmidsaurus.

### The Complete Genome collection

At several points in the manuscript, we refer to the Complete Genome collection. The Complete Genome collection includes the resulting assemblies from NCBI ONT, NCBI complete and Stanford Hospital *Enterococcus* isolate genomes. These genomes do not always reflect the definition of ‘complete’ assembly level from NCBI, yet they are all highly contiguous and taxonomically verified.

### IS counting

ISEScan (v1.7.2.3) was run on each genome with default parameters. Each resulting `{sample}.csv` file was used to count IS elements per sample, per contig, and per IS family.

### Assembly annotation

Genomes were annotated with bakta^35^ v1.9.1-1.10.3 using database v5.1 with the following flags

`--skip-tmrna --skip-ncrna --skip-ncrna-region --skip-crispr --skip-sorf --skip-gap --keep-contig-headers`. Output files including `.ffn`, `.faa`, `.gff3` were used for subsequent analyses.

### Analysis of IS transposase ORF distribution in *E. faecium*

For the identification and quantification of exact transposase open reading frame (ORF) copies in *E. faecium*, we analyzed the 602 *E. faecium* genomes from our NCBI complete and SH isolate collections combined (as described above). We excluded our NCBI ONT re-assembled genomes from this analysis because only 5 out of 175 datasets (Extended Fig. 1) were sequenced with the more recent R10.4.1 nanopore chemistry. Earlier-generation ONT data are known to exhibit higher rates of homopolymer errors^67^, which do not significantly hamper HMM-based approaches like ISEScan but would complicate accurate counting of exact copies of transposase nucleotide sequences.

Genome annotations were performed using bakta^35^, and the resulting GFF files were used to extract all CDS features longer than 500 nucleotides and labeled as “transposase”. These were then consolidated into a non-redundant transposase database, enabling us to quantify the frequency of each unique transposase nucleotide sequence across the dataset. We identified the six most abundant IS transposase families—ISL3, IS30, IS256, IS6, IS3, and IS110—and visualized their distribution (Fig. 1D). In some cases (due to available space on the plot) where bakta provided annotations more specific than the IS family level (e.g., “ISL3 family ISEfa11 transposase” vs. “ISL3 family transposase”), we retained these labels; choosing to include labels for high copy ISL3 elements, previously reported abundant IS elements in clinical *E. faecium* and IS1216, the IS6 element involved in the composite transposon (Tn1546) that confers vancomycin resistance.

The custom scripts for this analysis are available in the tpase_diversity_per_sample module of our GitHub repository.

### geNomad prediction of plasmid contigs

To predict whether a contig is likely from a chromosome or plasmid, we utilized geNomad^33^ (v1.8.0, v1.6 DB) with the `end-to-end` workflow. Contigs predicted as chromosomal with greater than a score of 0.6 were included in analyses as chromosomes. Contigs predicted as a plasmid with greater than a score of 0.7 were included in analyses as plasmid. Contigs not reaching those thresholds were considered undetermined and excluded from analyses where plasmid predictions were necessary (Fig. 1C, Extended Fig. 2).

### IS expansion in Enterococci

All *Enterococcus* genomes from the Complete collection and the 32 reference genomes described in Lebreton *et al*^36^, were phylogenetically classified with `de_novo_wf` from GTDB-Tk v2.4.0, Release 220^68^. In total, there were 2134 *Enterococcus* or outgroup genomes. As in Lebreton *et al.*, the term `g Vagococcus` was used to define the outgroup. A table summarizing the IS counts as defined by ISEScan for each genome, as well as the Newick and `tree.dist` files output by `de_novo_wf` were processed with the custom R script `is_expansion_phylogeny.R`. To visualize the ISL3 copy per genome among the *Enterococcus faecium* subclade, the genomes assigned to the representatives *EnGen0043*, *EnGen0015*, *Aus0004*, were added to a template for pruning the tree. A file containing the per-genome ISEScan values was prepared and uploaded to iToL^69^. To visualize counts per IS family across Enterococci, each genome was assigned to a single reference genome (1 of 32 reference genomes described in Lebreton *et al*^36^) based on the minimum distance to a reference. After assigning all genomes to a reference, the mean count of each IS family for the 32 sets of genomes was calculated and a file containing these values was uploaded to iToL.

### Defining infectious non-faecium non-faecalis *Enterococcus*

Non-*faecium* non-*faecalis* (NFF) *Enterococcus* causing human infection were based on the Australian *Enterococcus* Surveillance Outcome Program (AESOP)^37^. Over 10 years from 2013-2022, 11,681 *Enterococcal* bacteremia isolates were collected and identified. Any *Enterococcus* species responsible for at least 10 bacteremia isolates (average 1/year) in that collection is considered here as human-infectious. These species are *E. faecalis* (6,319), *E. faecium* (4,712), E. *casseliflavus* (198), *E. gallinarum* (166), *E. avium* (107), *E. raffinosus* (60), *E. hirae* (45), *E. lactis* (29), *E. durans* (27).

### Curation of short-read isolate data for ISL3 expansion timeline

A list of accessions was generated using the SRA search filter `((((Enterococcus faecium[Organism]) AND “illumina”[Platform]) AND “paired”[Layout]) AND “genomic”[Source]) AND “wgs”[Strategy]`, and metadata for these accessions was downloaded with the SRA Run Selector interface. Samples without a valid collection year or with estimated coverage (total bases / 3Mb) below 50x were removed. Samples from 2005 or later were filtered by location (Europe and North America) and source (any of `’Host’, ‘host_scientific_name’, ‘isolation_source’, ‘isolate_source’` metadata columns containing any of the terms `’blood’, ‘sapiens’, ‘patient’, ‘hospital’, ‘human’`), and then rarified to no more than 20 random samples per continent per year. Raw reads from the resulting 1559 samples were downloaded using SRA tools `prefetch` and `fasterq-dump`, deduplicated with `hts_SuperDeduper`, trimmed with `trim_galore`, and assembled with SPAdes. Assembly quality was assessed with QUAST, and taxonomy was verified with GTDB-Tk. Assemblies with inappropriate genome length (<2Mb, >3.4Mb), reference aligned length (<1.5Mb), or assigned species taxonomy were removed. 1378 assemblies passed filtering. The custom Nextflow workflow used to generate these results is available in our GitHub repository.

### IS counting in short read data

To quantify IS abundance in short-read assemblies, we first ran ISEScan on each assembly to identify IS families present on individual contigs. For each assembly, chromosomal coverage was approximated as the median coverage of contigs exceeding 5 kb in length. We then calculated a relative coverage for each contig by dividing the contig coverage by the chromosomal coverage. The number of IS copies on each contig was multiplied by its estimated coverage, and these values were subsequently summed across all contigs to obtain the final IS count per assembly.

We were concerned that changing IS abundance over time could be explained by recent isolates being disproportionately clinical in origin, while older isolates might derive from nonclinical sources, thus creating a spurious correlation. To control for this, we evaluated multiple IS families. Healthcare-associated *E. faecium* strains were previously reported to be enriched for IS256, IS30 and IS3 family elements. Consequently, if this kind of sampling bias were driving our findings, we would expect all IS families associated with clinical strains to follow the same temporal pattern (i.e., low in older and high in more recent samples). However, while IS256, IS30, and IS3 (linked to clinical clade A) are abundant even in older samples, ISL3 did not become prevalent until more recent time points. This finding indicates that the observed temporal trend in ISL3 abundance is unlikely to be an artifact of uneven sampling.

### Trendline of IS copy number over time

We used locally weighted scatterplot smoothing (LOWESS) to visualize temporal trends in per-genome IS counts. A smoothing fraction of 0.5 was chosen to balance local fit precision and global trend capture. Confidence intervals were generated via 1,000 bootstrap replicates. For each replicate, we randomly drew a subsample comprising 25% of the original data (with replacement) and applied LOWESS to these resampled points, evaluating the smoothed fit at each observed time. We pooled the smoothed fits across all bootstrap samples and calculated the 2.5th and 97.5th percentiles at each time point to outline a 95% confidence band.

### PopPUNK clustering of complete *E. faecalis* and *E. faecium* genomes for Pangraph input

To characterize structural variation among a set of nearly clonal genomes, we returned to the set of *Enterococcus* bloodstream infection isolates from Stanford Hospital as described above. With PopPUNK^43^ (v2.6.7), we clustered the 805 *E. faecium* Complete and Stanford Hospital genomes with the version 2 full *E. faecium* PopPUNK database downloaded from https://ftp.ebi.ac.uk/pub/databases/pp_dbs/Enterococcus_faecium_v2_full.tar.bz2. This was done with the `poppunk_assign` command with the additional flag of `run-qc`. 83 genomes were removed from clustering due to excessive core genome distances from the default type-isolate. PopPUNK’s `poppunk_visualize` command was used to generate a Newick file, `viz_core_NJ.nwk`, describing the resulting phylogeny. The output tree was visualized with iToL. We assigned sequence types with MLST^70^ (v2.23.0, available at https://github.com/tseemann/mlst) to each genome using the scheme `efaecium`. The Stanford hospital-endemic lineage corresponds to ST117 and formed a single cluster. The 68 genomes in the cluster, all from the Stanford collection, were selected for further analysis. Similarly, the 1,083 Complete and Stanford Hospital *E. faecalis* genomes were assigned to the version 2 full *E. faecalis* PopPUNK database downloaded from https://ftp.ebi.ac.uk/pub/databases/pp_dbs/Enterococcus_faecalis_v2_full.tar.bz2. None were removed by quality control. We attempted to assign sequence types as above using the scheme `efaecalis`. Here, the largest cluster corresponding to ST179 contained 57 genomes, 44 of which were Stanford isolates. The 44 genomes from the Stanford collection were selected for further analysis.

### Filtering genomes and stitching contigs for input into PanGraph

For each of the Stanford genomes in those clusters, we extracted the contigs predicted to be from the chromosome by geNomad as described before. Since Pangraph requires single contig inputs, we filtered for genomes with five or fewer contigs, yielding 66 *E. faecium* and 44 *E. faecalis* genomes. Multi-contig genomes were stitched together using 2kb ‘N’ spacers in the order output by Flye^71^.

### PanGraph analysis of structural variance in the SH *E. faecium* ST117 and *E. faecalis* ST179 clusters

For each of the two clusters defined above, we adapted the “Structural genome evolution” workflow as defined by Molari *et al*^47^. and found at https://github.com/mmolari/structural-evo.

Briefly, the genomes were processed with PanGraph^72^ (v0.7.1) to build a pangenome graph, which defines all “core blocks” (regions of shared homology across all genomes). Regions of structural variation are captured as edges between consecutive core blocks, and the set of sequences that can occur between two core blocks is defined as a “junction”. PanGraph was rerun on each junction (with its flanking core blocks), and the resulting “subgraph” defines each unique arrangement of homologous blocks through the junction as a unique “junction path”. Thus, for a given junction, the set of unique junction paths neatly describes the extent of structural sequence variation occurring in the cluster for a given structurally-variant locus.

Next, the set of junction subgraphs was analyzed with a custom Python workflow: first, we confirmed that the spacers (2kb ‘N’ as defined above) always occurred fully in the accessory region of junctions. This was expected, as disruptions in assembly contiguity are likely due to structural variation within the sample population itself. We removed any spacer-containing path from downstream analysis as their true sequence length and genotype is unclear. Next, for each junction, the workflow enumerates the distinct paths within the subgraph and characterizes the accessory genome of each non-empty path, using the bakta-derived annotations (described above) for each source genome. In ST117, the most common type of junction is a “binary” IS insertion. In that case, the junction comprises two paths, where one path is empty (i.e. there is no accessory region between the core blocks), and the other path contains only an IS element transposase annotation. IS-only junctions containing more than two paths and/or multiple IS families also occurred. Fewer large and more complex junctions occurred as well, including large insertions/deletions and phages, though we did not characterize these in detail.

As *E. faecalis* is the other ESKAPE *Enterococcus* species and has previously-reported IS abundance, the analysis of ST179 served as a control. The *E. faecalis* samples were collected from the same environment, location, and timeframe, and were processed and analyzed in an identical manner.

For compatibility with our high performance computing cluster, the PanGraph workflow was adapted to work for Snakemake v8.18.2. The workflow was run per cluster with existing installations of ISEScan (v1.7.2.3), DefenseFinder^73^ (1.3.0), geNomad (v1.8.0), and IntegronFinder^74^ (v2.0.5). The custom workflow was run with python 3.10 and polars v1.20, and is available in the pangraph module on our GitHub repository.

### Generation of circos plots of IS variants within PanGraph clusters

Code to generate the circos plot directly from the Pangraph result can be found in our GitHub repository. The isolate with earliest collection date, EF006 for the ST179 *E. faecalis* cluster and FM001 for the ST117 *E. faecium* cluster, was chosen as the reference coordinate system. The set of IS-only junctions was filtered to “backbone-only” junctions. In other words, IS insertions occurring in junctions not shared by all genomes (e.g. within other junctions or between core blocks that are inverted for some genomes) were not shown, as their position on the core backbone is ambiguous. Each junction was given a fixed width of 3 kb, as smaller sizes were not visible on the plot.

### ISL3 gene disruption and intergenic distance analysis in ST117

The *E. faecium* ST117 PanGraph analysis defined 100 ISL3 junctions (i.e., IS-only junctions containing only ISL3-family transposase annotation(s)) (Fig. 4D). To identify candidates for intragenic disruptions, we considered junctions where an annotated gene in the “empty” path (lacking the ISL3 insertion) overlapped fully or partially with the accessory region of a “filled” path. A junction was counted as gene-disruptive only if it contained such an overlapping gene and if that gene’s annotation was also missing or truncated in the filled path (Fig. 4E). The remaining ISL3 junctions were presumed to be intergenic. Of these, we considered every filled (ISL3-containing) path for every junction and extracted the distance from the 3’ end of the transposase to the next nearest gene, as well as its relative orientation. Paths that contained multiple accessory ISL3 transposases were only counted if they were all oriented the same way, otherwise the orientation was considered ambiguous and the path was excluded from analysis (Fig. 4F). As a control, the same analyses were done on the 51 IS30, IS256, IS110, IS3, and IS6 junctions as a combined set (“Other” in Fig. 4E and 4F).

### *E. faecium* domination longitudinal sample curation

To identify a set of metagenomic samples that were 1) clinically relevant, 2) longitudinal and 3) technically suitable for assembling complete *E. faecium* genomes directly, we selected longitudinal cases from our lab’s biobank of stool samples from individuals undergoing Hematopoietic cell transplantation (HCT) with high relative abundance of *E. faecium*. As previously described^49,53^, these stool samples were collected under an IRB protocol approved by The Stanford University Research Compliance Office (Protocol #8903; Principal Investigator: Dr. David Miklos, co-Investigator: Drs. Ami Bhatt and Tessa Andermann). Based on previous metagenomic sequencing results, samples with ≥10% relative abundance of *E. faecium* were considered. Any patients with at least two such samples were then identified and those samples ‘dominated’ (defined as ≥10% relative abundance) by *E. faecium* were selected for long-read metagenomic sequencing, yielding 34 samples from 14 patients (range 2-5 samples per patient).

### Metagenomic high molecular weight DNA extraction

To extract high molecular weight DNA from stool samples in preparation for metagenomic long-read sequencing, we used the QIAamp PowerFecal Pro DNA Kit (QIAGEN; 51804). Briefly, while working in a biosafety hood, from each stool sample kept on dry ice, ∼200 mg stool (2-3 biopsy punches) were added to the bead beating tube. Bead beating tubes were added in sets of 12 to a Vortex-Genie 2 with the horizontal Vortex Adapter described in the extraction kit manufacturer’s instructions and vortexed at max speed for 12 minutes at room temperature. All subsequent steps were consistent with manufacturer instructions using centrifugation. The final DNA was eluted in 100 µL C6 buffer and incubated at room temperature for 15 minutes before centrifugation. DNA concentration was measured by NanoDrop. DNA was then cleaned up with the Genomic DNA Clean&Concentrator-10 (Zymo Research; D4011) according to manufacturer instructions for large genomic DNA. Briefly, 200 µL of DNA binding buffer was added to 100 µL of DNA eluate from before. DNA was eluted in 25 µL nuclease free water. After clean-up, by Nanodrop, DNA concentrations ranged from 7.5-768.7 ng/µL per sample. Two samples had both DNA concentrations below 25 ng/µL and purity values (260/230) less than 1.5 and were not sequenced. DNA quality was confirmed with Genomic DNA ScreenTape Analysis (Agilent; 5067-5365, 5067-5366) on the Agilent 4150 TapeStation system. Samples had peak concentrations at DNA lengths greater than 20 kb. The loss of one sample, p11580_03 led to a singleton, but we continued with sequencing of p11580_06. Therefore 32 samples from 14 patients (range 1-4) were sequenced as described below.

### ONT sequencing, basecalling, quality control and assembly

Two pooled libraries containing 16 samples each were prepared with the Oxford Nanopore Native Barcoding 96-kit (SQK-NBD114.96) according to manufacturer’s instructions. Briefly, 400 ng of input DNA per sample was prepared, diluting the sample in nuclease free water if necessary. DNA concentration was confirmed for each sample with the Qubit DNA High Sensitivity kit at each checkpoint. Based on previous use of this approach for metagenomic sequencing of stool samples in our lab and TapeStation results, we expected our fragment length to be between 7-10 kb and therefore loaded 50 fmol of each library. Libraries were loaded onto separate R10.4.1 PromethION flow cells (FLO-PRO114M) according to manufacturer’s instructions and simultaneously sequenced on a PromethION 2 solo instrument. We anticipated assembling complete *E. faecium* genomes with at least 40x coverage. Therefore, we estimated that to reach 40x coverage of a 3 Mb *E. faecium* genome making up 10% relative abundance of the sample, we would need 1.2 Gb of sequencing per sample. To increase the likelihood of reaching this threshold for each sample and accounting for library imbalance, we let sequencing proceed until both libraries reached at least 2.5 Gb/sample of total sequencing. We obtained 60 Gb and 40 Gb of sequencing for each library, respectively.

Raw `.pod5` files were transferred to our computing cluster and basecalling was performed with Oxford Nanopore’s open source Dorado basecaller available at https://github.com/nanoporetech/dorado. The following command was used: dorado basecaller —kit-name SQK-NBD114-96 -b 1024 —trim ‘all’ sup@v5.0.0 {pod5} > {base called}.bam. Output bam files were demultiplexed and emitted as `.fastq` files with: `dorado basecaller —kit-name SQK-NBD114-96 -b 1024 —trim ‘demux’`. Of 32 sample barcodes, 31 had at least 1 Gb of sequencing after demultiplexing. One barcode corresponding to p11569_03 had less than 10 Mb of sequencing and was not considered further. Unfortunately, this made p11569_02 a singleton. Ultimately, 31 samples from 14 patients (range 1-4) were included for metagenomic assembly.

NanoPlot^75^ (v1.41.6) was used to evaluate sequencing quality per sample with default parameters. All 31 samples had median read quality of at least Q22.1 (22.1-24.2). Two samples, p11463_06 and p11580_06 had 9,150 and 11,220 reads respectively. Incidentally, these were the two samples where a single, long (>2.5 Mb), high coverage (100x), *E. faecium* contig could not be recovered (see below). The other 29 samples had at least 56,643 reads (56,643-659,377). The minimum read N50 for these 29 samples was 7.77 Kb. Metagenomic assembly was performed with Flye^76^ (v2.9.2) using the following command: `flye --nano-hq {reads} — meta —genome-size 200m —keep-haplotypes —no-alt-contigs`.

### Taxonomic profiling of assembled contigs with Sourmash

To identify *E. faecium* contigs from the metagenomic assemblies, all contigs 5kb or longer were taxonomically profiled with sourmash^77^ (v4.8.11) using the successive commands: `sourmash sketch dna -p k=51,abund –singleton {sample1}.fa` and `sourmash scripts fastmultigather -k 51 -m DNA –threshold-bp 5000 {samples}.zip gtdb-rs214-reps.k51.zip`. All contigs predicted as `g_Enterococcus;s_faecium` longer than 2.5 Mb, and with at least 100x coverage were considered ‘complete’ chromosomal contigs. One such contig was identified for 28/31 successfully sequenced samples as described above.

### Split *k-*mer analysis of *E. faecium* assemblies

To determine cases of ‘isogenic’ *E. faecium* chromosomes such that within strain heterogeneity and structural variation could be clearly assessed, the relatedness of the 28 *E. faecium* chromosomal contigs was considered with SKA^55^ (v1.0). All pairwise distances and clusters of highly related contigs were identified using the following commands: first, `ska alleles -f inputs.txt` and then, `ska distance -f skfs.txt -o results -S -s 40`, where `inputs.txt` contained paths to the 28 *E. faecium* chromosomal contig fasta files. Patients with longitudinal samples with < 10 SNPs/Mb or higher than 99.999% identity over time were considered for longitudinal within patient analyses. This threshold excluded p11342.

### Within-Sample and Within-Patient Structural Variant Calling

To assess within-sample *E. faecium* population heterogeneity, reads from each sample were aligned to their own metagenome assembly using ngmlr^78^ (v0.2.7) and default ONT sequencing presets. Structural variants (SVs) were called with Sniffles2^54^ (v2.6.0), which was run in both normal and mosaic modes to capture both dominant and subclonal variants. The outputs from these two modes were combined and any duplicate variants were removed, as well as any variants mapping to contigs other than the *E. faecium* chromosome and variants smaller than 100 nucleotides. The resulting per-sample SV counts are shown in Fig. 5B.

To determine whether ISL3 variants persisted or became fixed across sequential isolates from the same patient, we performed longitudinal variant calling: For each patient, the *E. faecium* chromosome from the sample collected at the earliest timepoint was chosen as the reference. Reads from each timepoint were mapped back to the reference metagenome assembly and passed as input to Sniffles2 as above. To look for SVs caused by IS element transpositions, all identified structural variants mapping to the *E. faecium* chromosome were screened to include only insertions or deletions in the 750–3000 bp size range, since this approximately corresponds to the length of IS elements. For each candidate variant, the predicted sequence was extracted and analyzed with ISEScan to confirm the presence of IS elements. The resulting IS variant calls were compiled, and patterns of persistence or fixation were examined by comparing the frequencies and presence/absence of these IS-containing variants across each patient’s timepoints. The custom scripts used for this analysis are available in the `lr_meta_sv` module of the GitHub repository.

### p11568 ISL3 structural variant neighborhood coverage

To measure the changing population frequency of the two identified ISL3 structural variants in the four sequenced p11568 samples over time, ONT reads from each of p11568 t1-t4 were aligned against the t4 *E. faecium* contig with minimap2^79^ with the flags `-x asm20`, `-c` and `-- eqx`. Per base read depth was then counted for each position with samtools^80^ (v1.21) using the command: `samtools depth -a -Q 10 -b t4.bed {sample}_sorted.bam > {sample}.coverage.txt`. Poor quality alignments are removed by the `-Q 10` flag. First, the mean alignment depth across the entire t4 *E. faecium* contig for each sample was calculated. Then, relative depth per base was calculated by dividing the depth at each position by the mean depth per sample. The t4 *E. faecium* contig was annotated with bakta as described above and the neighborhoods of each ISL3 structural variant was visualized.

### *E. faecium* isolation from stool

A single punch biopsy of stool samples from the first (t1) and last (t4) time points from p11568 were suspended in 300 µL PBS. Then, each sample was 10-fold serially diluted into 900 µL PBS to 1:10,000. 30 µL of the 1:10,000 dilutions were plated onto BHI agar and incubated overnight at 37°C. The next day, 6 colonies from each plate were selected and identified by biotyping with a MALDI-TOF based Bruker Biotyper per manufacturer instructions. Each identified *E. faecium* colony was streaked to isolation on a BHI agar plate. To confirm the insertion genotypes, overnights of isolates were prepared in 3 mL BHI broth and incubated with shaking at 37°C. Overnights were diluted 1:1000 into 10 mL subcultures and grown to OD ∼1.0. Cell pellets were then prepared according to Plasmidsaurus (Oregon, USA) ‘Standard Bacterial Genome with DNA extraction’ specifications. The resulting genomes were annotated with bakta as described above and the lack of both ISL3 insertions in t1, and the presence of both ISL3 insertions in t4 were confirmed in the assemblies.

### RNA-seq of *E. faecium* isolates

From glycerol stocks of both p11568 t1 and t4 confirmed to have the insertional genotypes (Fig. 6), four overnights were prepared in BHI as before. Each overnight was diluted 1:100 into 5 mL of BHI and subcultured for ∼3 hours until OD_600_ reached 0.4. 5 mL microtubes (AXYGEN, MCT-500-C) were prepared with 0.5 mL of quenching solution. The quenching solution contained 90% v/v 100% EtOH and 10% v/v saturated phenol for RNA extraction (ph 4-5). Microtubes were stored at -20°C until quenching occurred. After OD_600_ reached 0.4, 4 mL of each culture was transferred to a single prepared microtube. The microtube was vortexed until the solution was well-mixed. All microtubes were centrifuged at 10,000xg for five minutes at 4°C. The supernatant was poured off. The pellet was then lysed in 260 µL of a prepared mastermix containing 25:1 PBS:10mg/mL lysozyme (Sigma-Aldrich, L3790-10X1ML). Each pellet was resuspended in lysis buffer and then incubated at 37°C for 30 minutes while rocking on a BenchRocker. Then, 30 µL of 20% SDS was added followed by another round of incubation at 37°C. Then, 1.5 mL of Trizol was added and mixed-well by pipet. All subsequent steps were performed at room temperature. The solution was incubated for 10 minutes. Then 0.5 mL of chloroform was added and 10 vigorous, complete inversions of the tube were made before 3 minutes of incubation. After incubation, each tube was centrifuged at 12,000xg for 10 minutes. Carefully avoiding any non-aqueous layer, approximately 800 µL of the aqueous phase was transferred to a clean 2 mL eppendorf tube. Then an equal volume of 100% EtOH was added and mixed by pipet until emulsions were no longer visible. RNA was then purified using the Zymo RNA Clean & Concentrator-25 kit (R1018) according to manufacturer instructions. RNA was eluted in 25 µL nuclease-free water. RNA concentration was measured by NanoDrop and quality was confirmed by Bioanalyzer with the RNA 6000 Pico Kit (Agilent; 5067-1513). RNA was then stored at -80°C and sequenced with Novogene’s Prokaryotic Total RNA sequencing service (NovaSeq PE150, 3G raw data per sample).

### Analyses of RNA-seq data

Raw RNA-seq reads were trimmed with Trim Galore, available at https://github.com/FelixKrueger/TrimGalore/, using the command `trim_galore –quality 30 --length 60 --paired {sample}1.fq.gz {sample}_2.fq.gz -- retain_unpaired`. Bowtie2^81^ (v2.5.4) was used to align processed reads against the p11568 t4 isolate reference genome with default parameters. On the basis of a poor alignment fraction (20.99%) compared to the other replicates (minimum: 97.63%), the fourth replicate from the t1 isolate (t1_4) was excluded. To get the per base depth of reads, samtools depth was used as before except no quality threshold was implemented. This meant reads mapping equally well to multiple locations were distributed randomly. Furthermore, a feature count matrix filtering alignments with MAPQ < 10 was generated with bedtools (v2.27.1) multicov using the command `bedtools multicov -p -q 10 -bams {sample1}_sorted.bam {sample2}_sorted.bam …` against the t4 isolate reference genome. The resulting feature count matrix, along with the bakta annotation of the t4 isolate genome, were used to perform differential expression analysis with the DESeq2^82^ package (v1.42.1) in R. Only features with at least one count in both t1 (n=3) and t4 (n=4) were considered for differential expression. The adjusted p-value and log2(fold-change) for each of the 2,702 resulting features were visualized as a volcano plot. Thresholds of 1E-25 and |log2(FC)| ≥ 2 were used to define differentially expressed genes.

### *E. faecium* growth and antibiotic dose-response assays

*E. faecium* isolates were grown as described above. Antibiotic susceptibility testing was performed as described previously^56^, with some modifications. Briefly, each strain was grown overnight (∼18h) to stationary phase in BHI media at 37°C. Overnight cultures were then diluted to OD_600_=0.001 in Mueller-Hinton broth (MHB; Sigma-Aldrich) and incubated with two-fold serial dilutions of trimethoprim (TMP; Sigma-Aldrich) and sulfamethoxazole (SMX; Fisher) at the indicated concentrations. A ratio of 1:20 (TMP:SMX) was maintained for all treatments. Folic acid (Fisher) and folinic acid (Sigma-Aldrich) were supplemented at 1 µg/mL where indicated. Cultures were incubated at 37°C with continuous shaking in technical duplicate (150 μL volume per well) in a flat bottom 96-well microtiter plate (Corning) and sealed with a Breathe-Easy membrane (Diversified Biotech). Absorbance (600 nm) was monitored every 10 min for 24 hr using an Agilent BioTek Synergy H1 microplate reader. Growth was calculated as the percent area under the curve (AUC) relative to a control with no antibiotics added using the R package gcplyr^83^.

### Data visualization

Initial plots were generated by custom Python (Plotly) and R (ggplot) code. Plots were organized and refined with AffinityDesigner v2.5.7.

### Scripting

Custom bash, python, R, and Nextflow scripts referenced in the methods section are available in the GitHub repository. All scripts were run in R v4.3.2. Python scripts were run using v3.10 with polars v1.20.

## Data Availability

Long-read Enterococci genome assemblies, long-read metagenomic sequencing data, and RNA-sequencing data generated in this study are available under NCBI BioProject ID PRJNA1236482.

## Code Availability

Source code and custom workflows used for analyses in this manuscript are available on GitHub at https://github.com/abehr/ISL3-expansion-efaecium.

## End notes

### Acknowledgements

We are grateful to the patients, whose samples made this study possible. We thank Tessa Andermann and Andrew Rezvani for generating the Hematopoietic Cell Transplant patient stool collection and providing clinical expertise regarding the longitudinally sampled HCT patients. We would like to thank Sean Cuddihy, Angel Moreno, and members of the Stanford Clinical Microbiology Laboratory for their work isolating and banking *Enterococcus* from infection samples. We would like to thank Yewon Han of Martin Steinegger’s group for work on IS structural similarity. We would like to acknowledge the contributions of several members of the Bhatt lab for their assistance with experiments, Ashley Moore, Yishay Pinto and Jakob Wirbel. We would further like to acknowledge Jakob Wirbel, Rachael Chanin and Danica Schmidtke for their feedback on experimental design and manuscript preparation. This work used supercomputing resources provided by the Stanford Genetics Bioinformatics Service Center, supported by NIH S10 Instrumentation Grant S10OD023452. The Bhatt lab is supported by NIH R01AI148623 and R01AI143757, and a Stand Up 2 Cancer Grant. A.S.B. is supported by the Allen Distinguished Investigator Award. M.P.G is supported by Stanford’s Medical Scientist Training Program. A.A.B is supported by an NSF GRF award. This research was also supported, in part, by grant NSF PHY-2309135 to the Kavli Institute for Theoretical Physics (KITP). This paper describes the views of the authors and does not necessarily represent the official views of the National Institutes of Health (USA).

## Author Contributions

A.S.B, M.P.G and A.A.B conceived of and designed the study. M.P.G, A.A.B, S.B and J.D.L performed laboratory experiments. M.P.G, A.A.B, G.Z.M.R and E.F.B generated data from samples stored by N.B. Those samples were curated by M.P.G, G.R.N and J.L.S. M.P.G, G.Z.M.R and B.D curated the longitudinal HCT patient collection. Computational analysis was performed by A.A.B, M.P.G and M.M. The manuscript was drafted and all figures prepared by M.P.G and A.A.B. The manuscript was finalized by M.P.G, A.S.B and A.A.B.

## Competing Interests

The authors declare no competing interests.

## Additional Information

Supplementary Information is available for this paper.

## Extended Data

**Extended Data Fig. 1:**
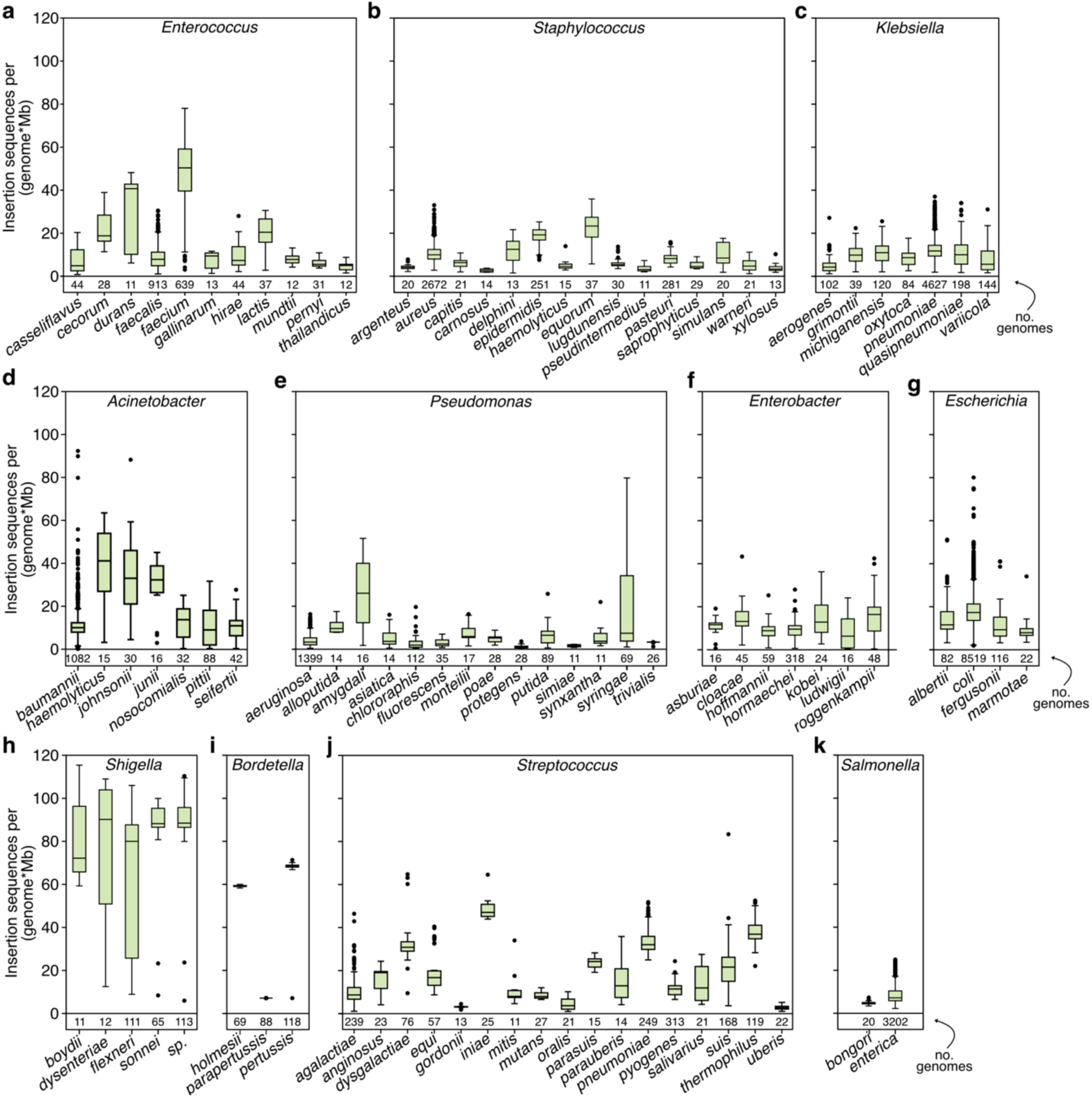
Total IS abundance across all surveyed species. Genomes from the NCBI ONT and NCBI complete collections passing quality filters as in Figure 1. Only species with at least 10 genomes defined by GTDB-Tk are included. The total IS counts per species are shown as a boxplot with the median, 25th and 75th percentiles, and individual values outside of 1.5 times the interquartile range (whiskers) visualized. Species are grouped by genus: **a**, *Enterococcus* **b**, *Staphylococcus* **c**, *Klebsiella* **d**, *Acinetobacter* **e**, *Pseudomonas* **f**, *Enterobacter* **g**, *Escherichia* **h**, *Shigella* **i**, *Bordetella **j**, Streptococcus* **k**, *Salmonella.* The y-axis scale is consistent across all panels and is based on the maximum across all species (*S. dysenteriae*). Several species of *Shigella* and *Bordetella* serve as positive controls for IS expansion.

**Extended Data Fig. 2:**
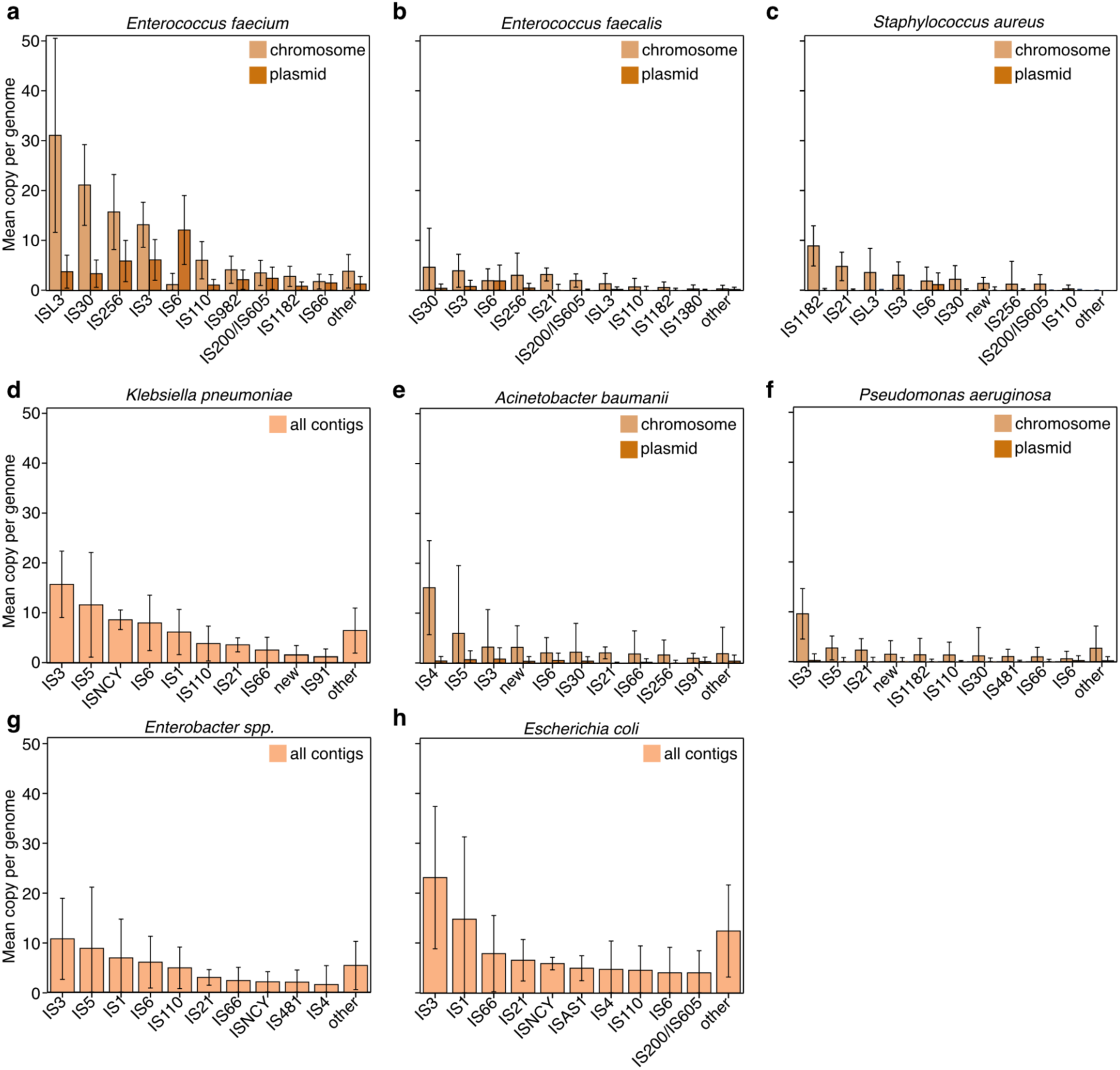
IS family abundance is surveyed in ESKAPE pathogen genomes. Mean (± s.d) insertion sequence copy per genome for each insertion sequence family as determined by ISEScan. These counts are not normalized by genome length. These data include ESKAPE genomes from the NCBI ONT and NCBI complete collections passing quality filters as in Figure 1. When possible, counts were assigned to chromosome or plasmid elements as defined by geNomad. For panels d, g, h, all counts are accumulated because geNomad predictions for genomes in these species were lower-confidence than for other species. **a**, *Enterococcus faecium* (same as Fig. 1C) **b**, *Enterococcus faecalis* **c**, *Staphylococcus aureus* **d**, *Klebsiella pneumoniae* **e**, *Acinetobacter baumanii* **f**, *Pseudomonas aeruginosa* **g**, *Enterobacter spp.* **h**, *Escherichia coli.* The y-axis scale is consistent across all panels and is based on the maximum across all species (ISL3 in *E. faecium*).

**Extended Data Fig. 3:**
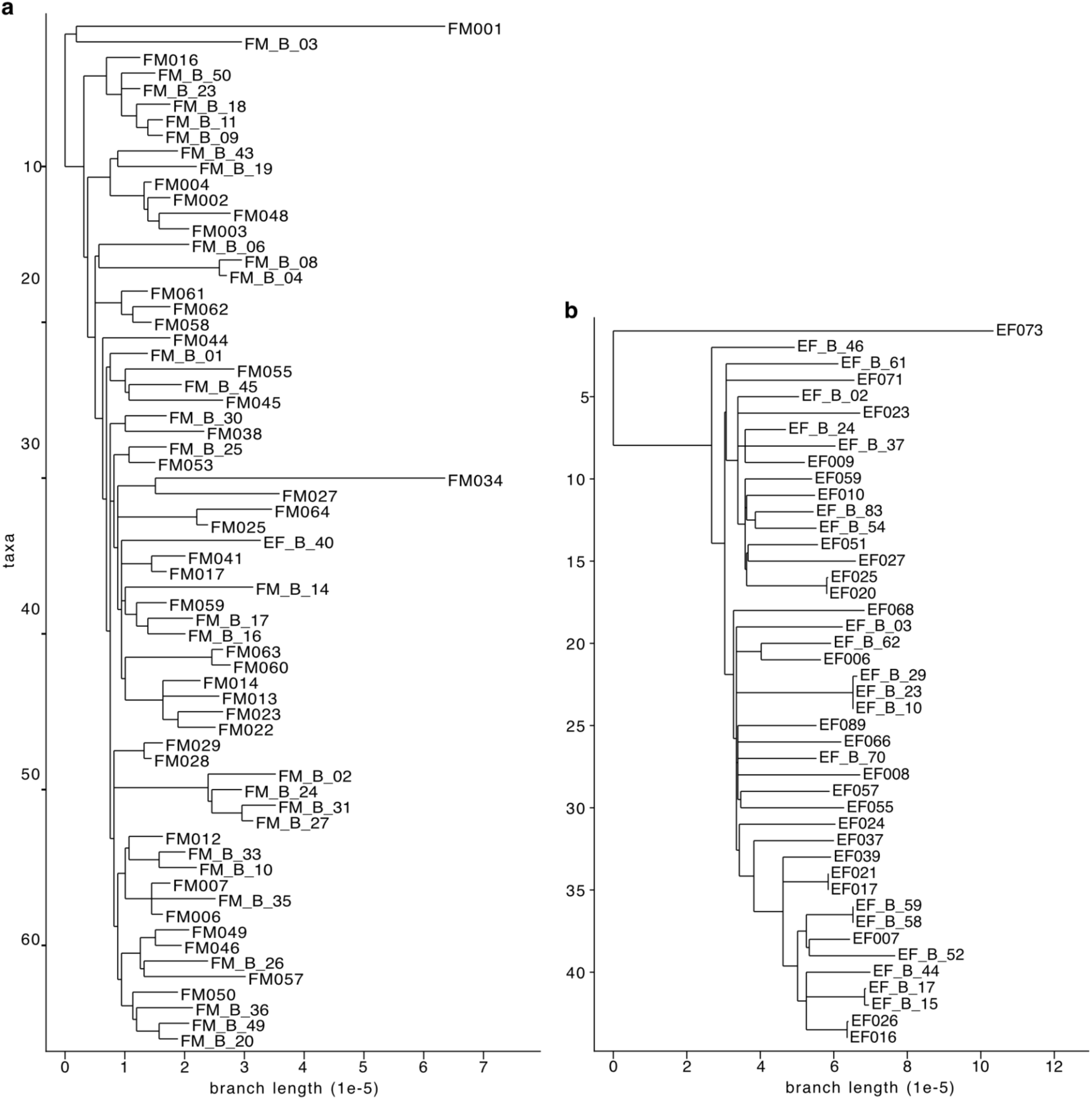
Pangraph input cluster core genome phylogenetic trees after filtering recombination. **a**, 66 *E. faecium* genomes making up the Stanford Hospitals ST117 cluster. The polished alignment after filtering out putative recombination regions was 1.587 Mb and the tree was constructed from 865 SNPs. **b**, 44 *E. faecalis* genomes making up the Stanford Hospitals ST179 cluster. The polished alignment after filtering out putative recombination regions was 2.522 Mb and the tree was constructed from 2,355 SNPs. **a**, **b** Branch length is equivalent to the number of SNPs per core-genome alignment length.

**Extended Data Fig. 4:**
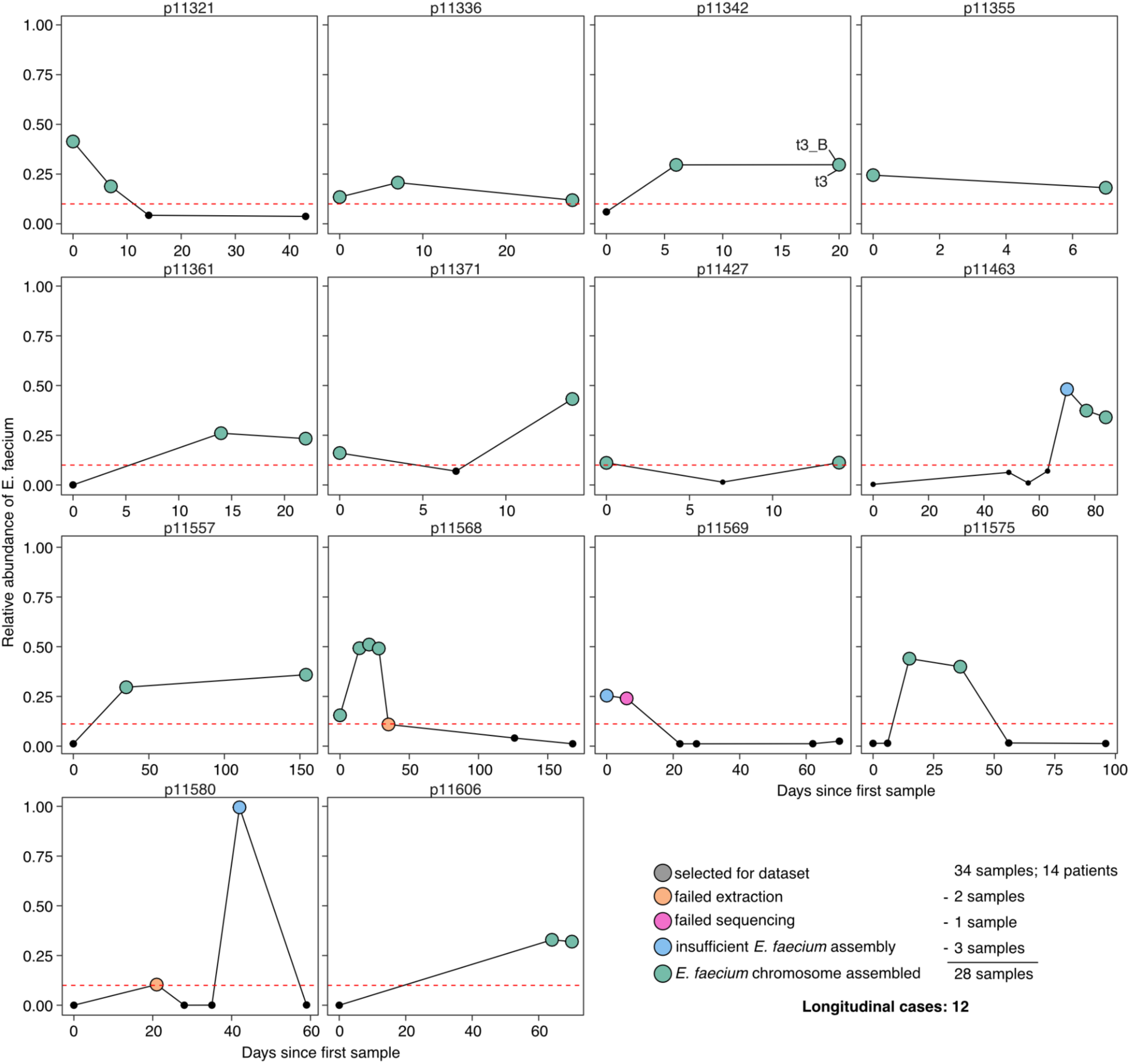
Longitudinal Hematopoietic Cell Transplant (HCT) patient cohort for *in situ* structural variant detection. The relative abundance of *E. faecium* as measured by previous short- or linked-read metagenomics of these samples for each of 14 potential longitudinal cases^53^. Among the 34 samples from 14 patients, *E. faecium* chromosomes could not be assembled due to failed extraction (2 samples), failed sequencing (1 sample) or insufficient assembly (3 samples) for 6 samples. The resulting 28 *E. faecium* chromosomes comprise 12 longitudinal cases. Note, for p11342, t3 was sampled twice and both were successfully assembled.

**Extended Data Fig. 5:**
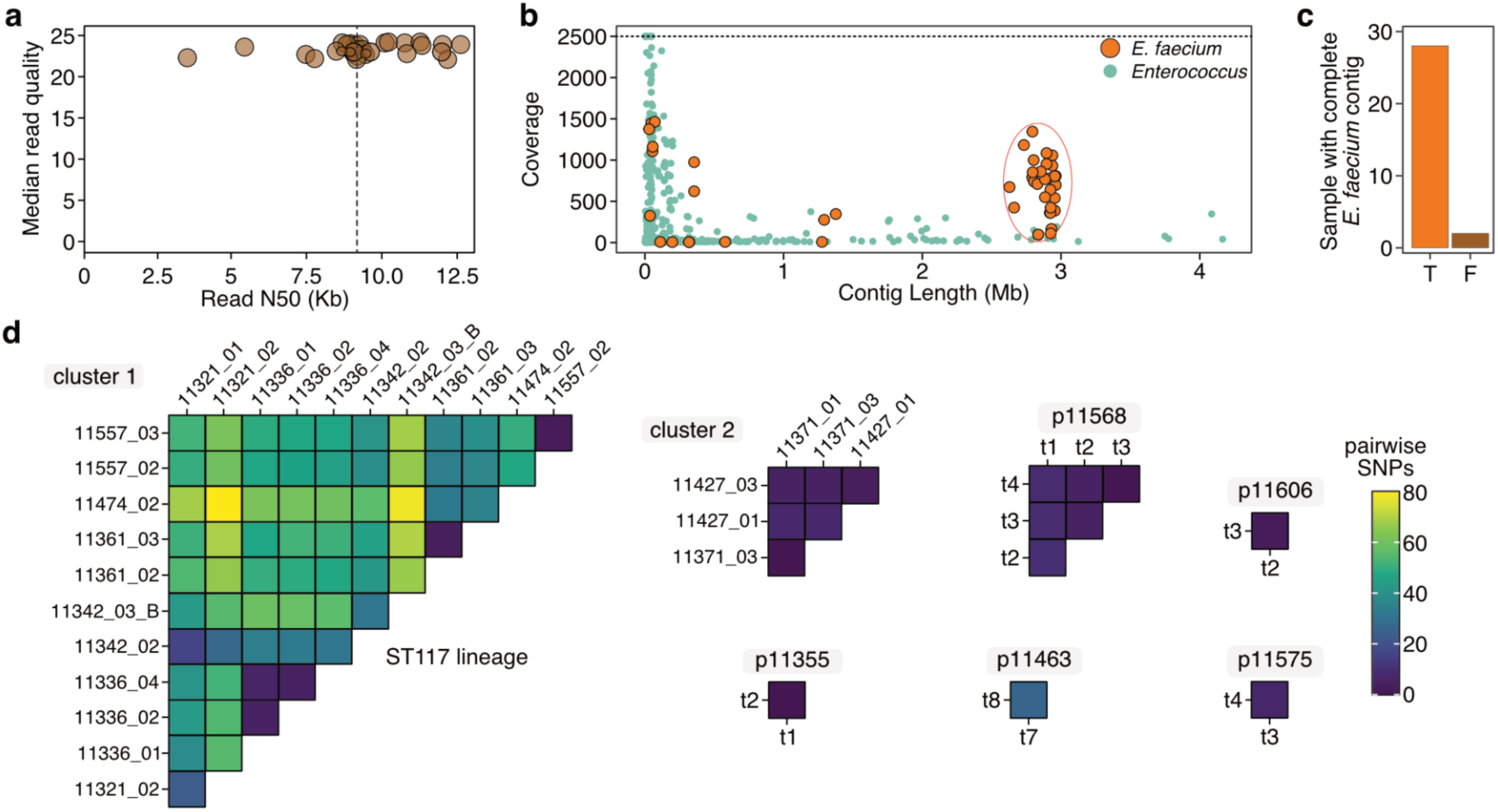
Longitudinal long-read metagenomic sequencing, assembly and verification of strain identity over time. **a**, Extracted DNA was prepared for long-read sequencing with the Oxford Nanopore Native Barcoding 96-kit (SQK-NBD114.96) and sequenced on an R10.4.1 flow cell. Median read quality was >Q22 for all samples and the median Read N50 is ∼9.2 Kb (dashed line). **b**, Extracted contigs longer than 5kb were taxonomically profiled with Sourmash. Coverage and contig length (Mb) for those contigs predicted to be within g_*Enterococcus* are plotted. A group of *E. faecium* contigs with at least 100x coverage and longer than 2.5 Mb were considered as *E. faecium* chromosomes for further analysis (orange oval). Coverage was elided at 2500x (dashed line) **c**, 28/31 samples input for sequencing had an *E. faecium* chromosome. **d**, All pairwise comparisons of *E. faecium* chromosome identity as measured by SKA. Clusters were generated with a SNP threshold of 40. Cluster 1 and cluster 2 contain multiple patients while the other 5 clusters contain samples from a single patient.

**Extended Data Fig. 6:**
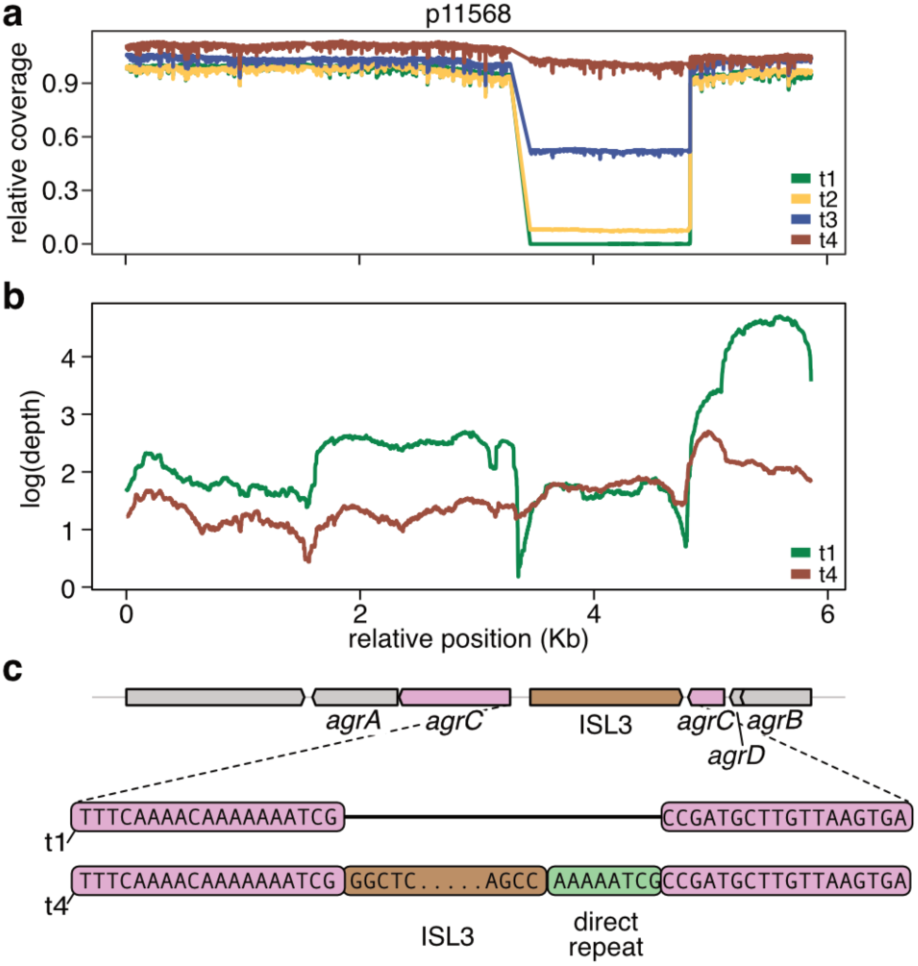
ISL3 disruption of *agrC* silences the *agr* operon. **a**, In the neighborhood of the ISL3 variant fixing within the *agr* operon, the relative coverage of long-reads aligned to the metagenomic assembled *E. faecium* chromosome of p11568 t4, normalized to the average chromosomal coverage per sample. The coordinates are relative to the locus (Kbp) **b**, From *E. faecium* isolated from p11568 t1 and t4 stool samples, RNA-seq read depth (log-scale) across the same locus as in panel a. **c**, The sequence context of the ISL3 structural variant within the *agr* operon is examined in detail. The position of the ISL3 insertion (brown), the direct repeat (green), and the split *agrC* CDS (purple) are highlighted.

**Extended Data Fig. 7:**
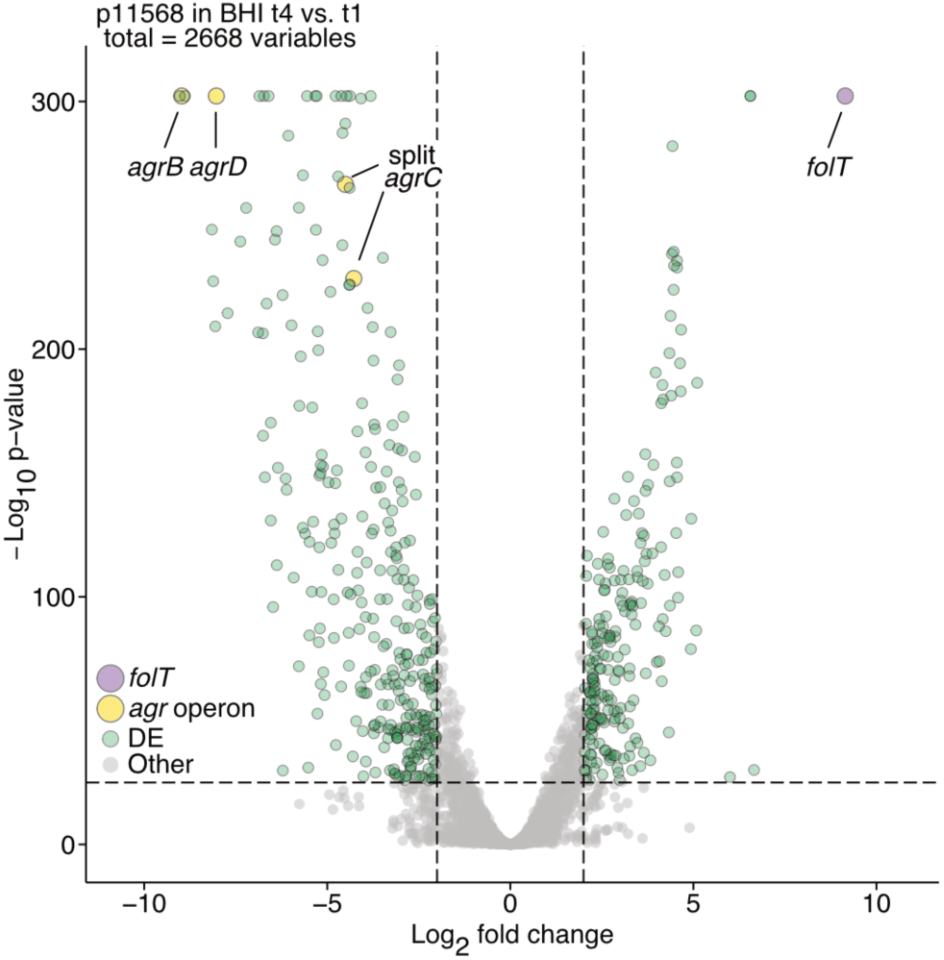
ISL3 variants have profound effects on neighboring gene expression. A volcano plot showing the relative gene expression of the isolates from t4 vs. t1 (t1: n=3, t4: n=4). Only alignments with MAPQ ≥ 10, and features having at least one count in a replicate of both t1 and t4 were considered (n=2,668). An adjusted p-value and the fold change (log2-scale) of genes were calculated by DESeq2. DE genes (green) and the *agr* operon (yellow, enlarged), the *folT* gene (purple, enlarged), were additionally labeled.

**Extended Data Fig. 8:**
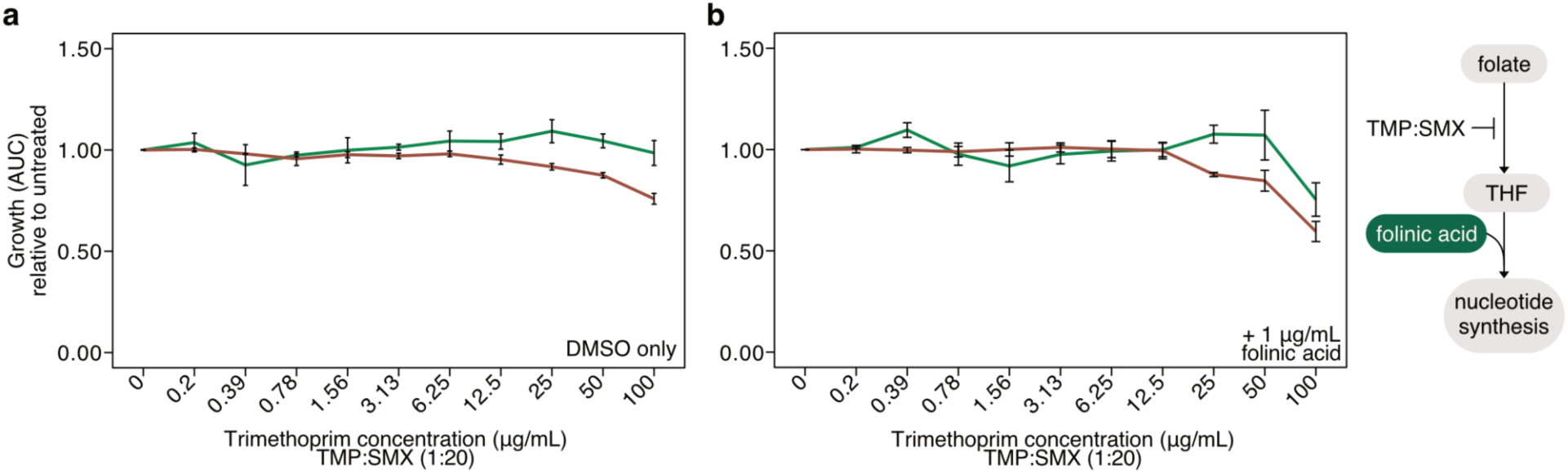
An activated folate metabolite rescues growth of p11568 t1 and t4 *E. faecium*. **a**, To measure the effect of drug solvent alone, relative to untreated, the growth (area under the curve, AUC) of the t1 and t4 isolates in increasing concentrations of dimethylsulfoxide (DMSO) at the equivalent drug concentrations (concentration of TMP is labeled, TMP:SMX was prepared in a ratio 1:20). **b**, Relative to untreated, the growth (area under the curve, AUC) of the t1 and t4 isolates in increasing concentrations of TMP:SMX (concentration of TMP is labeled, TMP:SMX was prepared in a ratio 1:20), with all conditions supplemented with 1 µg/mL folinic acid. (right) A simplified schematic showing where TMP:SMX and folinic acid act within the folate activation pathway.

## Supplementary Information

**Supplementary Data 1**

IS count table: All publicly available genomes, annotated with their source collection, taxonomy (NCBI or GTDB-Tk depending on collection), and counts for each IS family.

**Supplementary Data 2**

IS counts for SH *E. faecium* and the several SH atypical Enterococci genomes generated in this study.

**Supplementary Data 3**

PanGraph junction path statistics for the *E. faecium* ST117 cluster (worksheet 1) and the *E. faecalis* ST179 cluster (worksheet 2).

**Supplementary Data 4**

Sequencing statistics (nanotstats, worksheet 1) and assembly statistics (flye contig info, worksheet 2) for the long-read metagenomic samples generated in this study.

**Supplementary Data 5**

Modified Sniffles output in VCF-like format from longitudinal metagenomic data showing structural variants within each sample (worksheet 1) and longitudinally across samples per patient (worksheet 2).

**Supplementary Data 6**

RNA-seq data: DESeq results (worksheet 1) and feature count matrix (worksheet 2).

**Supplementary Data 7**

Set of *E. faecium* genomes with ISL3 proximal to *folT*, along with their relative position and orientation.

## Notes

### Competing Interest Statement

The authors have declared no competing interest.

